# Sphingosine-1-Phosphate Receptor 1 regulates competition dependent astrocyte morphogenesis and tiling in murine cortex

**DOI:** 10.64898/2026.03.28.714989

**Authors:** Jean Patrick M. Gonzales, Connor Tuck, Surya Chandra Rao Thummu, Soha Munir, Jaden Harris, Hunain (Zeik) Tariq, Paul Marcelli, Oscar Dominguez, Kaviya Suriyakumar Anbazagan, Sandeep K. Singh

## Abstract

Astrocytes are highly abundant in the mammalian brain and coordinate with neurons and other glial cells to regulate neural circuit structure, function, and blood brain barrier integrity among many to maintain proper brain homeostasis. Astrocytes perform most of these functions owing to their highly complex morphologies and hundreds of thousands of fine processes that are important in contacting neuronal synapses and other glial cells. In fact, the morphological complexity of astrocytes is regulated by the presence and activity of neurons and helps establish astrocyte territory/tiling in a non-overlapping pattern; however, the mechanisms of astrocyte tiling are not well characterized. Using a human astrocyte-mouse neuron coculture system, we previously showed that sphingosine-1-phosphate receptor 1 (S1PR1) regulates astrocyte morphogenesis in a neuronal contact dependent manner. In this study, we find that S1PR1, *in vivo,* regulates astrocyte morphogenesis in a cortical layer specific manner. Using astrocyte-specific S1PR1 knock out mouse models and adenoassociated viral labeling methods, we show that S1PR1 is crucial in establishing competition driven astrocyte tiling and morphogenesis in the developing brain. Furthermore, we show that JAK-STAT3 signaling regulates neuronal contact induced expression of S1PR1 in cocultured astrocytes. These studies therefore uncover a lipid signaling receptor as a major regulator of astrocyte morphogenesis and tiling in murine cortical layers.

## Introduction

Development of the central nervous system (CNS) is stringently coordinated and regulated by intricate physical, biochemical, and spatiotemporal modes of cross-communication between nervous system cells. Within the brain, cells form dynamic and functional neural-glial networks through the integration of cell-cell interactions, growth factor signaling, and neural activity, which culminate in the activation of neurogenic, downstream intracellular signaling cascades. These interconnected networks provide the structural and functional foundation for higher-order brain processes—including cognition, learning and memory, perception, and emotion—facilitating swift processing of information to adapt to changes in environmental stimuli. Improper assembly of neural-glial networks is a characteristic feature of numerous neurodevelopmental and neurological disorders, including Autism and Alzheimer’s disease, highlighting the importance of understanding the key players and molecular mechanisms that govern this process^1–3^.

Astrocytes are one of the key players participating in neural-glial networks and are a versatile cell type in the mammalian brain^4,5^ Defined by their highly polarized and elaborate morphology, astrocytes extend ramified processes into the surrounding parenchyma to directly contact or come into proximity with neighboring neurons, glia, and vasculature^6^. These processes serve as an interface for intercellular communication through which astrocytes perform a range of essential functions: they provide metabolic and trophic support to energy-demanding neurons^7^, coordinate with microglia to remove cellular debris and prune excess synapses during development^8,9^, facilitate oligodendrocyte-mediated myelination of axons^10^, and establish non-overlapping domains by maintaining boundaries with adjacent astrocytes^11^. Although the molecular cues and intrinsic genetic programs governing astrocyte differentiation are well characterized^12,13^, the intercellular signaling events that guide astrocyte maturation and sculpt their morphological complexity during postnatal brain development are not fully understood.

While protein-centric effectors and signaling pathways have been extensively studied in the context of astrocyte differentiation and development^14,15^, emerging evidence highlights the critical contribution of lipids—particularly sphingolipids—in astrocyte biology. Sphingolipids are a diverse class of bioactive lipids that are synthesized *de novo* through conserved, endoplasmic reticulum-resident metabolic pathways and are enriched in the brain, where they serve both structural and signaling roles^16^. In cellular membranes, sphingolipids help organize lipids into microdomains, or “lipid rafts,” which concentrate signaling complexes and regulate membrane trafficking^17,18^. More importantly, many sphingolipid metabolites act as potent intracellular and extracellular signaling molecules that influence cell survival, proliferation, differentiation, and morphology^19^. Within the CNS, sphingolipids have well-documented roles in neuronal development and synaptic transmission^20,21^; however, their functions in astrocytes remain less defined. Recent studies suggest that sphingolipid metabolites, such as Sphingosine-1-Phosphate (S1P), regulate astrocyte proliferation, migration, and morphological differentiation^22–24^. S1P, acting through its G-protein-coupled receptors, can influence cytoskeletal dynamics to promote process extension, a hallmark of astrocyte morphogenesis^25–27^. In addition, altered levels of S1P and genes within the S1P metabolic pathway are altered in many neuropathologies, including Parkinson’s disease, Alzheimer’s disease, and neuropathic pain; yet the fundamental role of the S1P-S1PR1 axis in the brain remains less explored.

Previously, we’ve shown that the expression of the G-protein-coupled receptor S1PR1 increases during postnatal brain development in mice and is primarily localized to perisynaptic astrocytic processes (PAPs) and regulates astrocyte morphogenesis in a neuronal contact-dependent manner. Furthermore, exogenous S1P led to a marked increase in astrocyte complexity and expression of genes encoding astrocyte-secreted synaptogenic factors (SPARCL1 and TSP4) when cocultured with neurons. These findings suggest that the S1P–S1PR1 signaling axis modulates neuron-astrocyte interactions in promoting astrocyte morphological maturation and their synapse association^28^. More recently, *in vivo* studies using *drosophila* (invertebrate) and zebrafish (vertebrate) models have further substantiated the role of S1P–S1PR1 signaling in astrocyte morphogenesis. The astrocyte-specific deletion of S1PR1 in zebrafish, or its functional homolog Tre1 in Drosophila, resulted in significantly reduced astrocyte volume and branching complexity^29^. However, the role of S1P-S1PR1 signaling in higher-order, mammalian astrocytes has yet to be elucidated.

In this study, we investigated the role of S1PR1 in regulating astrocyte morphogenesis during murine postnatal brain development. Our findings demonstrate that S1PR1 modulates murine astrocyte size, branching complexity, and territorial tiling in a cortical layer-specific manner. In addition, our *in vitro* astrocyte-neuron coculture experiments identified the JAK/STAT3 signaling pathway as a regulator of neuronal contact-dependent expression of S1PR1 in astrocytes. Together, these findings position S1P-S1PR1 signaling as a novel regulator of murine astrocyte development in a local neuronal signaling-driven manner, underscoring its importance in establishing the complex morphology required for astrocyte function and overall brain homeostasis.

## Materials and Methods

### Mice

All mice used in this study were maintained on a C57BL/6J background. Knockout mice were generated by crossing S1PR1^loxP/loxP^ (a gift from Dr. Laura Sim-Selley, VCU) mice with GFAP-Cre mice (JAX, #024098). To label astrocytes and their processes, we intracranially injected either AAV pZac2.1-GfaABC1D-YFP, AAV pZac2.1-GfABC1D-Lck-GFP, AAV pZac2.1-GfABC1D-YFP-P2A-Cre, AAV pZac2.1-GfaABC1D-tdTomato, or AAV pZac2.1-GfaABC1D-tdTomato-P2A-Cre in P2-P3 pups. Four weeks after injection, the mouse brains were collected and S1PR1 expression was confirmed via immunofluorescence staining. Afterwards, astrocyte morphological imaging and analysis was performed. To determine the cell-type specificity of the GFAP-Cre mouse line, we crossed unfloxed GFAP-Cre mice with the B6J.129(B6N)-Gt(ROSA)26Sor^tm1(CAG-cas9*,-EGFP)Fezh/^J (or Rosa26 loxP-STOP-loxP Cas9-P2A-GFP mouse (JAX # 021675)) to generate a Cre-dependent reporter mouse. In this line, Cas9 and GFP expression is limited to cells expressing Cre thereby allowing to report specificity and efficiency of GFAP-Cre promoter. For *in vitro* experiments, P1 and P3 animals were used for neuronal and astrocyte cultures, respectively. The above experiments were approved by Virginia Commonwealth University IACUC.

### Primary neuron isolation

Mouse cortical neurons were isolated from wild-type C57BL/6 P1 pups as described earlier^30^. Briefly, the cortices were dissected and digested with papain (Worthington Biochemical Corp) in the presence of DNase at 36^°^C for 45 mins (with intermittent swirling every 15 mins), followed by trituration in low and high concentrations of ovomucoid. The cells were passed through 40 µm nitex filters (Fisher Scientific) and subjected to a series of negative panning plates coated with 2× Bandeiraea Simplicifolia Lectin I (Vector Laboratories) followed by incubation at room temperature (RT) in AffiniPure goat anti-mouse IgG + IgM (H + L) (Jackson Immunoresearch Laboratories, PA) and anti-neural cell adhesion molecule L1 (MAB5272, Millipore) coated positive panning dishes to obtain >95% pure cortical neurons. Following incubation, the cells were collected, centrifuged at 1100 rpm for 15 mins, and resuspended in neurobasal growth medium (NGM) comprised of neurobasal medium, B27, 100 U/mL Pen/Strep, 2 mM L-Glutamine, and 1 mM Na+Pyruvate. All the media components were procured from Gibco unless otherwise stated. The cells were counted and seeded onto poly-D-lysine (PDL)/laminin-coated coverslips at a density of 2 x 10^5^ cells per coverslip in a 24-well plate and incubated in a humidified chamber with 10% CO₂ at 37°C. AraC (1 uM) was added onto the wells for the initial 24-36 h to prevent the growth of glial cells. Every 3 days, half of the neurobasal medium was replaced with fresh medium until the day of the experiments.

### Astrocyte isolation from S1PR1-GFP mouse brain

*S1pr1^eGFP/eGFP^* (B6.129P2-S1pr1tm1Hrose/J)(JAX # 028623) mouse cortical astrocytes were obtained from newborn pups (P3) as described earlier^30^ with slight modifications. Briefly, the cortices from the pups were dissected and the tissue was digested with papain in the presence of DNase at 36° C for 45 minutes. Papain-digested tissue was triturated with low and high ovomucoid solutions, filtered through nitex mesh (20 µm), and resuspended in serum free neurobasal growth medium (exactly as was for neurons). The cells were counted using an automated cell counter, and 10-14 million cells were plated in 75 cm^2^ flasks coated with poly-D-lysine and incubated at 37°C in 10% CO_2_. On DIV-3, neurobasal medium was removed and rinsed with DPBS (-MgCl_2_ and -CaCl_2_) and shaken by hand for 10–15 seconds or until a monolayer of astrocytes remained. The cells were rinsed once with DPBS (-MgCl_2_ and -CaCl_2_) and replenished with fresh medium. Upon confluency (typically around DIV6-7), astrocytes were trypsinized with 0.25% trypsin and passaged to 6-well dishes with 7-8 x 10^5^ cells per well and maintained till around DIV10.

### Neuron-astrocyte coculture experiments

For neuron-astrocyte co-culture experiments, the astrocytes cultured from S1PR1-GFP pups at around DIV8-DIV10 were detached with 0.25% trypsin following which trypsin was neutralized using DMEM containing 10% FBS medium. The cells were triturated to create a single-cell suspension, centrifuged at 200g for 11 minutes, and resuspended in neuronal growth medium; cell counts were taken, and ∼3 x 10^4^ cells were added to each well of a 24 well plate containing WT neurons. Separate wells were maintained where only astrocytes were added to the coverslips coated with PDL and laminin that served as astrocyte monoculture/control. Following incubation in the presence and absence of neurons, the cells were fixed with 4% PFA.

### Inhibitor treatment of neuron-astrocyte co-cultures

For experiments investigating the role of signaling pathways such as JAK-STAT, Sonic Hedgehog, and S1P-S1PR1 interaction in regulating astrocyte morphology and S1PR1 expression, astrocyte monocultures or neuron-astrocyte co-cultures were treated with different inhibitors such Ruxolitinib (1 µM), Sonidegib (5 µM), and W146 (200 nM), respectively. Inhibitors or vehicle (DMSO) control was added to respective wells 30 min. prior to adding astrocytes into the wells with or without DIV7-9 neurons. In tandem with co-culture experiments, astrocyte monocultures were treated with inhibitors along a similar timeline. Twenty-four-hour post incubation, cells were fixed, immunostained, and imaged for analysis.

### Immunofluorescence analysis of frozen brain tissues

Mice were anesthetized using isoflurane inhalation and transcardially perfused with 1× PBS pH 7.4 followed by 4% paraformaldehyde. Brains were harvested, fixed in 4% paraformaldehyde overnight, and placed in 30% sucrose for 48-72 h before being embedded in a 2:1 30% sucrose:OCT solution. Coronal brain slices of 50 µm (morphological analyses using YFP or Lck-GFP labelling) and 20 µm (immunofluorescence analyses) were cryosectioned on a cryostat, rehydrated in 1× PBS for 15 min, blocked with 10% goat serum in PBS with 0.5% Triton x-100 for 1 hour, primary antibody incubation overnight at 4^°^C, followed by a 2 hour secondary antibody incubation at RT, and mounted using VECTASHIELD Antifade Mounting Media with DAPI (Vector Laboratories, H-1000). The following primary antibodies were used: Mouse anti-GFAP (1:500; Cell Signaling Technology, 3670S), Rat anti-CD31 (1:200; Abcam, ab56299), Rabbit anti-AQP4 (1:500; Alomone, AQP-014), Rabbit anti-S1PR1 (1:100; Santa Cruz, sc-48356), Guinea pig anti-GLT1 (1:500; EMD Millipore, AB1783), Rabbit anti-NeuN (1:500; Cell Signaling #24307), Rabbit Iba1 (1:1000; Wako Corporation #019-19741), Chicken anti-OLIG2 (1:500; Antibodies Incorporated # OLIG2-0100), Rabbit anti-SOX9 (1:500; Cell Signaling; #82630) and Chicken anti-GFP (1:1000; Aves; # GFP-1010). The following secondary antibodies were used: Alexa Fluor 594 goat anti-rat (1:200; Thermo Fisher Scientific, A11007), Alexa Fluor 488 goat anti-rabbit (1:500; Thermo Fisher Scientific, A11034), Alexa Fluor 488 goat anti-Mouse (1:500; Thermo Fischer Scientific, A32723), Alexa Fluor 594 goat anti-Guinea pig (1:500; Thermo Fischer Scientific, A11076), and Alexa Fluor 488 goat anti-Chicken (1:500; Thermo Fisher Scientific #A32931TR).

### Immunofluorescence analysis of astrocyte-neuron cocultures

For immunostaining, cells were fixed in 4% paraformaldehyde (PFA) at room temperature for 10 minutes. Following fixation, cells were rinsed with 1x PBS twice at room temperature (RT), followed by blocking with blocking buffer containing 50% Normal Goat Serum (NGS) and 0.2% Triton-X100 in antibody solution. Cells were then incubated in antibody solution with 10% NGS overnight at 4°C with the following antibodies: anti-GFP (1:2000, Aves; # GFP-1010), anti-S100b (1:500, Millipore) and anti-Map2 (1:2000, Cell Signaling; #8707). Subsequently, the cells were washed 3x with 1X PBS and incubated with species-specific secondary antibodies conjugated with Alexa Fluor 488 (1:500) or Alexa Fluor 594 (1:500) in an antibody solution with 10% NGS for 1h at RT. Coverslips were then washed three times with 1X PBS and mounted with VECTASHIELD Antifade Mounting Media (Vector Laboratories, H-1000). Cells were imaged at 20x magnification using a confocal microscope (Zeiss LSM 710).

### Confocal Imaging and Image analysis

For analysis of astrocyte morphology *in vitro*, GFP signals from astrocytes were used for measuring complexity using the Sholl analysis plugin in FIJI. Briefly, images were smoothened (process-smooth) and converted to a binary signal. The freehand tool was used to generate a region of interest (ROI) around the astrocyte and remove processes from neighboring cells. Using the straight, segmented, or freehand lines tool, an ROI was marked at the center of the cell and stretched to the longest processes of the astrocyte, followed by execution of Sholl analysis (Plugins-neuroanatomy-Sholl-Sholl analysis). For measurement of the intensity, area, and perimeter of individual astrocytes, the green channel was converted to a grayscale image, followed by threshold adjustment, which was set the same for all the images. Using the wand tracing tool, each cell was selected and measured. To evaluate astrocyte morphology *in vivo*, fluorescence images were acquired using the Zeiss LSM 880 laser-scanning confocal microscope with a ×40 oil-immersion objective with a frame size of 1,024 × 1,024 and bit depth of 16 (Zen3.1). Serial images at the z-axis were taken at an optical step of 0.5 um, with the overall *z*-axis range encompassing entire astrocytes. Images were imported into Imaris Bitplane software, and only astrocytes with their soma between the *z*-axis range were chosen for further analysis. We performed 3D surface rendering using the Imaris Surface module to characterize astrocyte volume and surface area. Morphological analysis was performed using the Imaris Filament module. Astrocyte branches and processes were outlined by Autopath with the starting point set at 8 μm and the seed point set at 0.5 μm, and statistical outputs, including ‘filament number Sholl intersections,’ were extracted and plotted with Prism software. To analyze the knockout efficiency of *S1PR1*, fluorescence images were acquired using a Zeiss LSM 880 laser-scanning confocal microscope with a 20x objective. To measure the fluorescence intensity of GFAP and AQP4 fluorescence images were acquired using the Keyence BZ-X1000 fluorescence microscope with a 20x objective and analyzed using Fiji. GLT1 images were acquired at 63X magnification in Zeiss 880 confocal microscope and analyzed using Fiji. Brains from the GFAP-Cre Cas9 reporter mice were stained and imaged for NeuN, Iba1, OLIG2, GFAP and GFP using a Zeiss LSM 880 laser-scanning confocal microscope with a 20x objective. The person who analyzed the images was blinded to the experimental groups.

### Tracer dye experiment to measure Blood-Brain Barrier Integrity

Brain tracer leakage experiments were performed as described^27,28^. Mice (4-5 wk of age) were injected in the tail vein with Alexa Fluor 555–cadaverine (6 μg/g; Invitrogen) or 3-kDa dextran–TMR (10 μg/g; Invitrogen) dissolved in saline. After 2 h (for Alexa Fluor 555–cadaverine or 3-kDa dextran–TMR) mice were anesthetized and perfused for 7 min with ice-cold PBS (pH 7.4), and brains were removed. After dissection, the cortex was weighed and homogenized with 1% Triton X-100 in PBS. Cortical lysates were centrifuged at 12,000 × *g* for 20 min at 4°C, and the supernatant was used to quantify fluorescence (excitation/emission 540/590 nm; SpectraMax M2e; Molecular Devices). The relative fluorescence values were normalized with cortical weights.

### AAV packaging

All AAVs used in this study were produced in the Singh lab unless otherwise mentioned, in accordance with the US National Institutes of Health Guidelines for Research Involving Recombinant DNA Molecules and the Virginia Commonwealth University Institutional Biosafety Committee. pAAV-GfaBC1D-YFP-P2A-Cre plasmid was created by subcloning P2A-Cre from pAAV-hSynapsin-mCherry-P2A-Cre-WPRE plasmid (a gift from Hui Yang (Addgene plasmid # 107312); into the pAAV-GfaBC1D-YFP plasmid. Similarly, pAAV-GfaBC1D-tdTomato and pAAV-GfaBC1D-tdTomato-P2A-Cre was created by subcloning and replacing YFP with that of tdTomato into pAAV-GfaBC1D-YFP or pAAV-GfaBC1D-YFP-P2A-Cre plasmid respectively. The serotype AAV plasmid, PHP.eB (pUCmini-iCAP-PHP.eB), was a gift from Viviana Gradinaru (Addgene plasmid #103005; RRID: Addgene_103005). pAAV.GfaBC1D-PI-Lck-GFP-SV40 was a gift from Baljit Khakh (Addgene viral prep # 105598-AAV5).

AAV packaging was performed as previously described^31^. Briefly, ∼1x10^8^ HEK293T cells (ATCC #CRL 3216) grown in 6, T-175 flasks were transfected using polyethylenimine (PEI) with the pAAV of choice, the helper plasmid pAd-ΔF6 (Addgene Plasmid (#112867), and the serotype plasmid AAV PHP.eB at a ratio of 1:1:2 using polyethylenimine transfection. The media was changed the following day, and cells were cultured for an additional 48 hours. After 72 hours post-transfection, cells were scraped, pelleted, resuspended in cell lysis buffer (15mM NaCl, 5mM Tris-HCl, pH 8.5), followed by 3, 10-minute freeze-thaw in a dry ice/ethanol bath followed by a 37° C water bath. Cell lysates were treated and incubated at 37°C with benzonase (50 U/ml) for 30 minutes and centrifuged at 4500 rcf for 30 min at 4℃ to collect the supernatant. The supernatant was added to a 15%, 25%, 40%, and 60% iodixanol gradient loaded into a polypropylene ultracentrifuge tube (∼1% phenol red was incorporated into the 25% and 60% layer to visualize the density gradient), and underwent ultracentrifugation at 67,000 rpm using a Beckman Ti-70 rotor for 1 hour at 18 °C. After ultracentrifugation, the AAV fraction was collected using a syringe with an 18 G needle and concentrated with four washes of DPBS (Millipore, 100 K MW) via centrifugation at 4000 × *g* for 10 minutes each. A final volume of 200 μl of virus after the final buffer exchange was recovered from the column, aliquoted and stored at -80°C.

### Intracranial injection of AAV viruses

For astrocyte labeling, we utilized pAAV-GfaABC1D-YFP, pAAV-GfaBC1D-Lck-GFP, pAAV.GfaBC1D-YFP-P2A-Cre (sparse KO), pAAV-GfaBC1D-tdTomato, and pAAV-GfaBC1D-tdTomato-P2A-Cre (Territory overlap). In brief, pups aged post-natal days 2-3 were anesthetized on ice, followed by a 0.8 µL viral injection (∼1x 10^9^ virus particles) using a Hamilton neurosyringe microinjector at a depth of 0.5 mm in the right hemisphere of the visual cortex. After injections, pups were gently placed on a heating pad to recover and closely monitored. All of the animal procedures were performed in accordance with approved VCU IACUC protocols.

### Astrocyte territory overlap/tiling analysis

P3 S1PR1^f/f^ pups were intracranially injected with 1 μL of a combination of control AAVs (GfaABC1D-YFP + GfaABC1D-tdTomato) or Cre AAVs (GfaABC1D-YFP-P2A-Cre + GfaABC1D-tdTomato-P2A-Cre) mixed at a 1:1 ratio, respectively. At P30, the mouse brains were collected, cryosectioned, and 50 µm Z-stack (0.5 µm Z-intervals) images which contained two neighboring astrocytes (expressing either YFP or tdTomato) were collected. Astrocytes containing endfeet were excluded from this experiment.

IMARIS 11 was used to generate a 3D surface reconstruction of the YFP and tdTomato astrocyte to generate fluorescent channel masks of each astrocyte. Filament traces were then generated from each astrocyte to utilize the “Convex Hull” Imaris Xtension to generate a surface representing the territory of each astrocyte. Territory overlap was measured by creating a mask of the original fluorescent channel within the convex hull, and the colocalization tool in Imaris was used to generate a colocalization channel representing territory overlap between the masked YFP and tdTomato channels. Then, a surface of the colocalization fluorescent channel was generated to record the volume of territory overlap. Territory overlap volume measurements were normalized to the territory volume of both the YFP (green) and tdTomato (magenta) astrocyte to control for variations in the astrocyte size, as the 50 µm Z-stack limitation prevented capture of at least one astrocyte in its entirety.

### Quantification and statistical analysis

Sample sizes and statistical tests can be found in the accompanying figure legends. Offline analysis was performed using Graphpad Prism 11 and Microsoft Excel. Statistical significance was determined by unpaired 2-tailed Welch’s t-tests for comparison of 2 groups. Data are presented as mean ± s.e.m. Statistical significance is indicated by asterisks: \**P* < 0.05, \*\**P* < 0.01, \*\*\**P* < 0.001, \*\*\*\**P* < 0.0001. The number of independent experiments (*n*) and the number of analyzed cells is noted in the figure legends.

## Results

### Astrocyte morphology and S1PR1 expression are driven by neuronal contact *in vitro*

We have previously demonstrated that the expression of S1PR1 in the developing mouse brain coincides with synaptogenesis and the maturation of astrocytes, and that neuronal contact stimulates S1PR1 expression in astrocytes^28^. To further investigate S1PR1’s expression and localization dynamics, we prepared primary astrocyte cultures from S1PR1 reporter mice (S1PR1^GFP/GFP^) (JAX Strain # 028623) in serum-free medium. It is well established that astrocytes cultured in the presence of serum can adopt a reactive phenotype and exhibit altered morphology and gene expression^32^. To avoid these confounding effects, our astrocytes were maintained in serum-free Neurobasal medium, and we verified the cellular purity of these cultures (Fig. S1 and data not shown). We validated S1PR1 expression by probing for GFP in astrocytes marked with S100b and found that S1PR1 signal primarily localizes to astrocytic plasma membranes (Fig. S1B). Therefore, we can simultaneously measure S1PR1 expression (quantified as GFP fluorescent intensity) and astrocyte morphological parameters (perimeter and Sholl analyses) using GFP signal. S1PR1-GFP astrocytes cultured alone exhibit a simple, flattened fibroblast-like morphology and basal S1PR1 signal. Interestingly, most astrocytes cocultured with neurons exhibit increased S1PR1 expression, three-dimensionality, and extensive branching (Fig. S1B,C); however, some astrocytes still maintained a simple morphological state with low S1PR1 signal (Fig. S1C). To determine whether neuronal contact induces S1PR1-GFP expression and drives astrocyte complexity, S1PR1-GFP astrocyte/WT neuron cocultures were immunostained with the neuronal marker, MAP2 (Fig. 1A,E). This approach allowed us to identify neuronal processes and explicitly measure astrocytes in direct contact (termed interacting astrocytes, or IA) versus no contact with neurons (termed non-interacting astrocytes, or NIA) (Fig. 1A,E). Indeed, S1PR1-GFP astrocytes in direct contact with MAP2^+^ neurons exhibited much higher S1PR1-GFP fluorescent intensity and branching, as indicated by Sholl analyses (Fig. 1D,H) and increased perimeter (Fig. 1C,F) compared to astrocytes with no neuronal contact. Time course experiments also demonstrate that the neuronal contact-dependent increase in S1PR1 expression and astrocyte complexity already begins to show at 8 hours (Fig. 1A-D) and increases further at 24 hours (Fig. 1E-H). Therefore, we provide evidence that neuronal contact directly and simultaneously induces S1PR1 expression and astrocyte morphological complexity. To further test the role of S1PR1 signaling in contact-dependent astrocyte morphological complexity, we applied the S1PR1-specific antagonist, W146^31^ in these astrocyte-neuron cocultures and analyzed the astrocyte only conditions or IAs (Fig. 1I,K). As expected, W146 significantly reduced neuronal contact-dependent astrocyte complexity as measured by a reduction in perimeter (Fig. 1J) and number of Sholl intersections (Fig. 1L), affirming the role of S1P-S1PR1 signaling in astrocyte morphogenesis. To our surprise we also found that W146 significantly reduced neuronal-contact-induced expression of S1PR1 in astrocytes (fig. 1K) but had no effect on astrocyte only cultures (Fig. 1K), further suggesting S1PR1 signaling may also self-regulate its expression in astrocytes when in contact with neuron.

**Fig 1.**
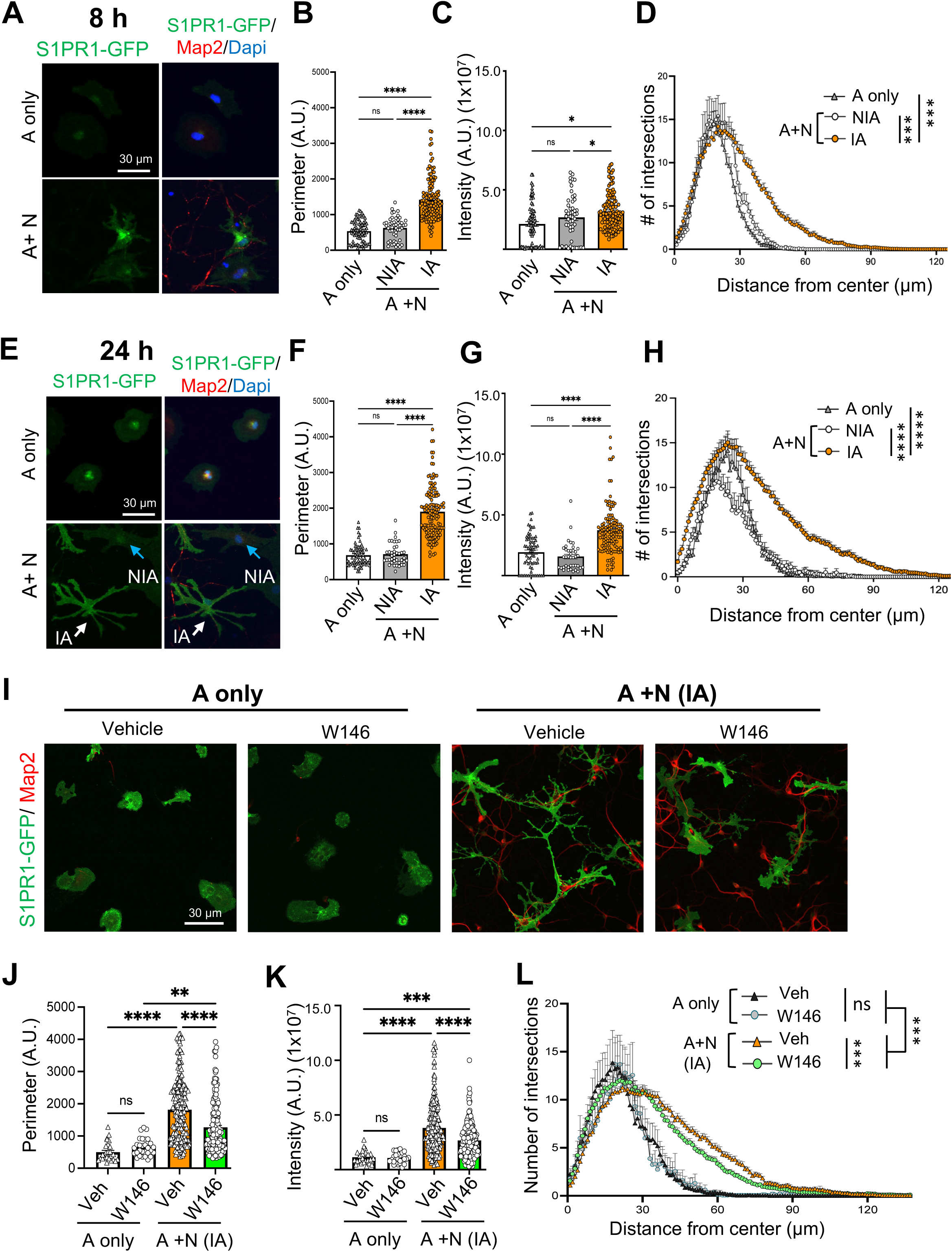
Astrocyte morphology and S1PR1 expression is driven by neuronal contact *in vitro*. Representative confocal images (20X) of S1PR1-GFP astrocytes (green) cultured alone (**A only**) or with WT neurons (**A+N**) stained with Map2 (red) for 8 **(A-D)** or 24 hours (**E-H**). Based on the astrocytes interaction with or without Map2+ neuronal processes in A+N cocultures, astrocytes were separated into either non-interacting astrocytes (NIA) (Blue arrow in E) or interacting astrocytes (IA) (white arrow in E). Quantification of perimeter (**B** and **F**) and S1PR1 expression as green signal intensity (**C** and **G**) per astrocyte, respectively, in A or A+N cultures. (**D** and **H**) Quantification of astrocyte morphological complexity as indicated by number of intersections plotted against distance from center of the soma by Sholl analyses from A or A+N cultures. (**I-L**) S1PR1 inhibition blocks neuronal contact dependent astrocyte morphological complexity. (**I**) Representative confocal image of S1PR1-GFP astrocytes treated with vehicle (DMSO) or the S1PR1 antagonist, W146, for 24 hours cultured with or without neurons (Map2, red). Perimeter (**J**) and fluorescent intensity (**K**) analyses of GFP immunofluorescence from S1PR1-GFP astrocytes. (**L**) Quantification of morphological complexity of astrocytes as indicated by number of intersections plotted against distance from center of the soma by Sholl analyses. Data represents the mean ± SEM from two- (A-H) or three (I-L) -biological replicates. * = *p*<0.05, *** = *p* < 0.001, **** = *p*<0.0001; one way ANOVA.

Next, we sought to identify signals that induce expression of S1PR1 in astrocytes and thus in turn regulate astrocyte morphogenesis. Several signaling pathways, including JAK-STAT3 and SHH-GLI1, have been shown to play crucial roles in neuron-astrocyte interactions and modulate astrocyte maturation and morphological complexity^32–34^. Therefore, we applied chemical inhibitors of these pathways, Ruxolitinib and Sonidegib, respectively, to our coculture paradigm (Fig. 2). Interestingly, while Ruxolitinib and Sonidegib both inhibited astrocyte morphological complexity (Fig. 2B,D,E), neuronal contact induced expression of S1PR1 was significantly inhibited by only Ruxolitinib in cocultures (Fig. 2C). These results demonstrate JAK-STAT3 signaling but not SHH signaling mediates neuronal contact-dependent expression of S1PR1 in cocultured astrocytes.

**Fig 2.**
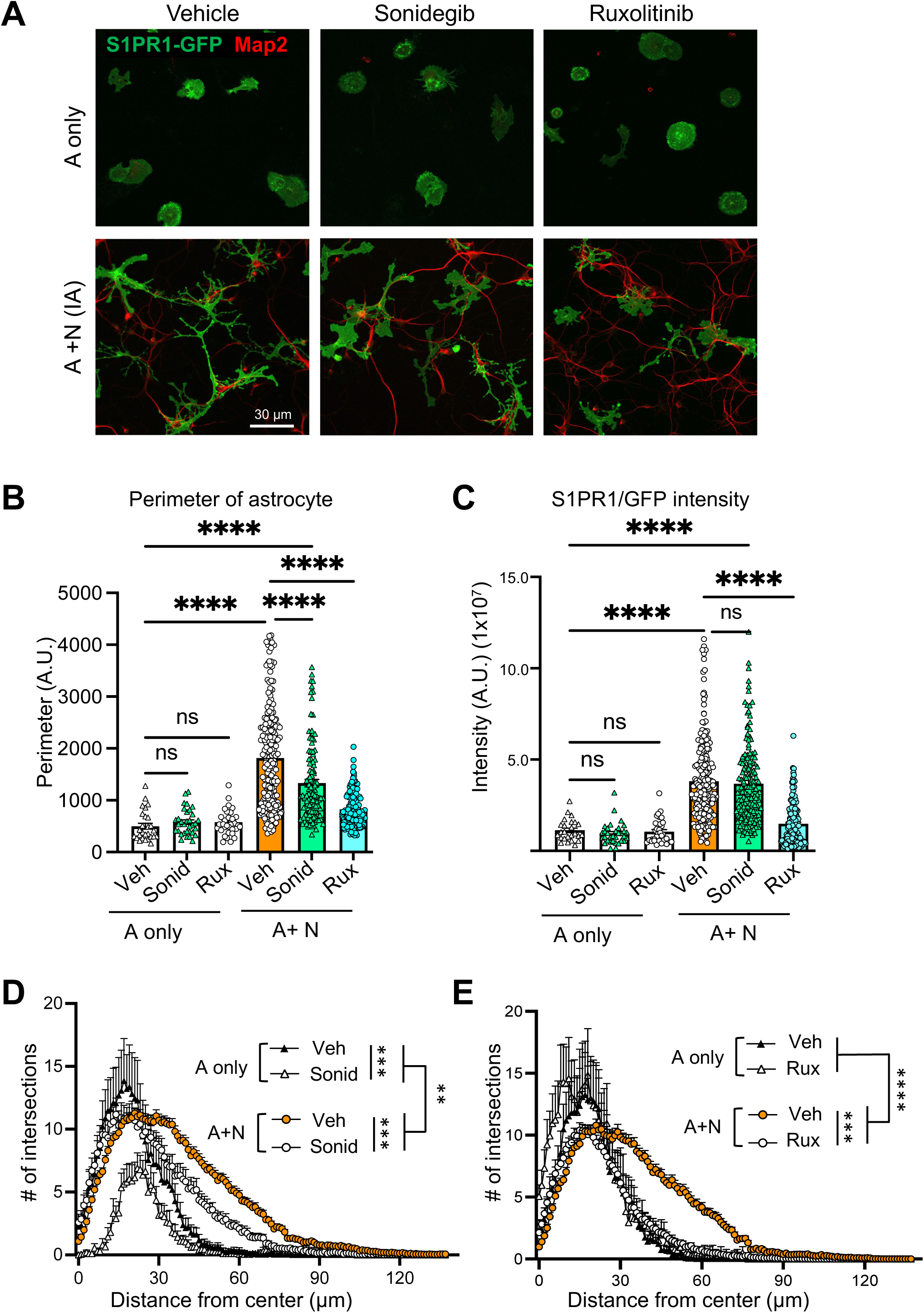
JAK/STAT but not Sonic hedgehog signaling regulates neuronal contact-induced S1PR1 expression in astrocytes. **(A)** Representative confocal images (20X) of S1PR1-GFP astrocytes (green) cultured alone (**A only**) or with WT neurons (**A+N**) stained with Map2 (red) treated with vehicle (DMSO), the JAK2 inhibitor Ruxolitinib (Rux), or the Smoothened inhibitor Sonidegib (Sonid), for 24 hours. Quantification of perimeter (**B**) and level of S1PR1 expression as fluorescent signal intensity (**C**) per astrocyte respectively in A or A+N cultures. (**D, E**) Quantification of astrocyte morphological complexity as indicated by the number of intersections plotted against distance from center of the soma by Sholl analyses from A or A+N cultures treated with vehicle (DMSO) control, Rux, or Sonid. Data represents the mean ± SEM from three biological replicates. ** = *p*<0.01, *** = *p* < 0.001, **** = *p*<0.0001, one way ANOVA.

### Astrocyte-specific S1PR1 knock-out mice exhibit no gross changes in blood vasculature, astrocyte maturation and astrocyte numbers

To characterize the role of astrocytic S1PR1 in the developing brain *in vivo*, we generated an astrocyte-specific S1PR1 knockout mouse line (termed S1PR1^ΔAST^) in which the S1PR1 gene is exclusively deleted in astrocytes by crossing S1PR1^loxP/loxP^ mice with the GFAP-Cre (77.6) mouse line (Fig. 3A). S1PR1 expression at the mRNA level was drastically reduced in the cortices of S1PR1^ΔAST^ mice at P30 compared to littermate controls (Fig. 3B), demonstrating astrocytes as the major source of S1PR1 in the brain *in vivo*. We further performed immunofluorescence staining and confocal imaging of S1PR1 in the cortices of S1PR1^ΔAST^ and littermate controls and found strong expression of S1PR1 within the cortex of control brains and a drastically reduced signal in S1PR1^ΔAST^ cortices (Fig. 3C). Interestingly, endothelial S1PR1 expression remained unchanged in S1PR1^ΔAST^, as evidenced by colocalization of the endothelial cell marker, CD31, with S1PR1 (Fig. 3C, arrow). This observation is in line with the well-known expression of S1PR1 in endothelial cells and is crucial for vascularization and maintenance of blood-brain barrier (BBB) integrity^35^. Because S1PR1 is expressed on astrocytic endfeet^28^ and astrocyte endfeet are crucial in maintaining BBB integrity, we investigated whether astrocytic S1PR1 may also participate in BBB integrity and astrocyte-vasculature interactions. We immunostained S1PR1^ΔAST^or littermate control brains at P30 with the astrocyte endfeet marker, AQP4, and endothelial cell marker CD31 (Fig. 3D,E) and observed no significant changes in the signal corresponding to the area and fluorescent intensity of both AQP4 and CD31, implicating that astrocytic S1PR1 does not affect the general structure of vasculature and endfeet (Fig. 2C-E). To conclusively demonstrate astrocytic S1PR1’s role in the functional maintenance of BBB integrity, we performed dye tracer leakage experiments, which are well-established methods to measure BBB leakage^35^. In this assay, we intravenously injected two tracer dyes, either a 3-kDa dextran or the smaller 1-kDa cadaverine, and measured their relative accumulation in the brain tissue (Fig. 3F). Leakage of both tracer dyes was comparable between cohorts (Fig. 3F), suggesting that astrocytic S1PR1 does not play a prominent role in BBB integrity.

**Fig 3.**
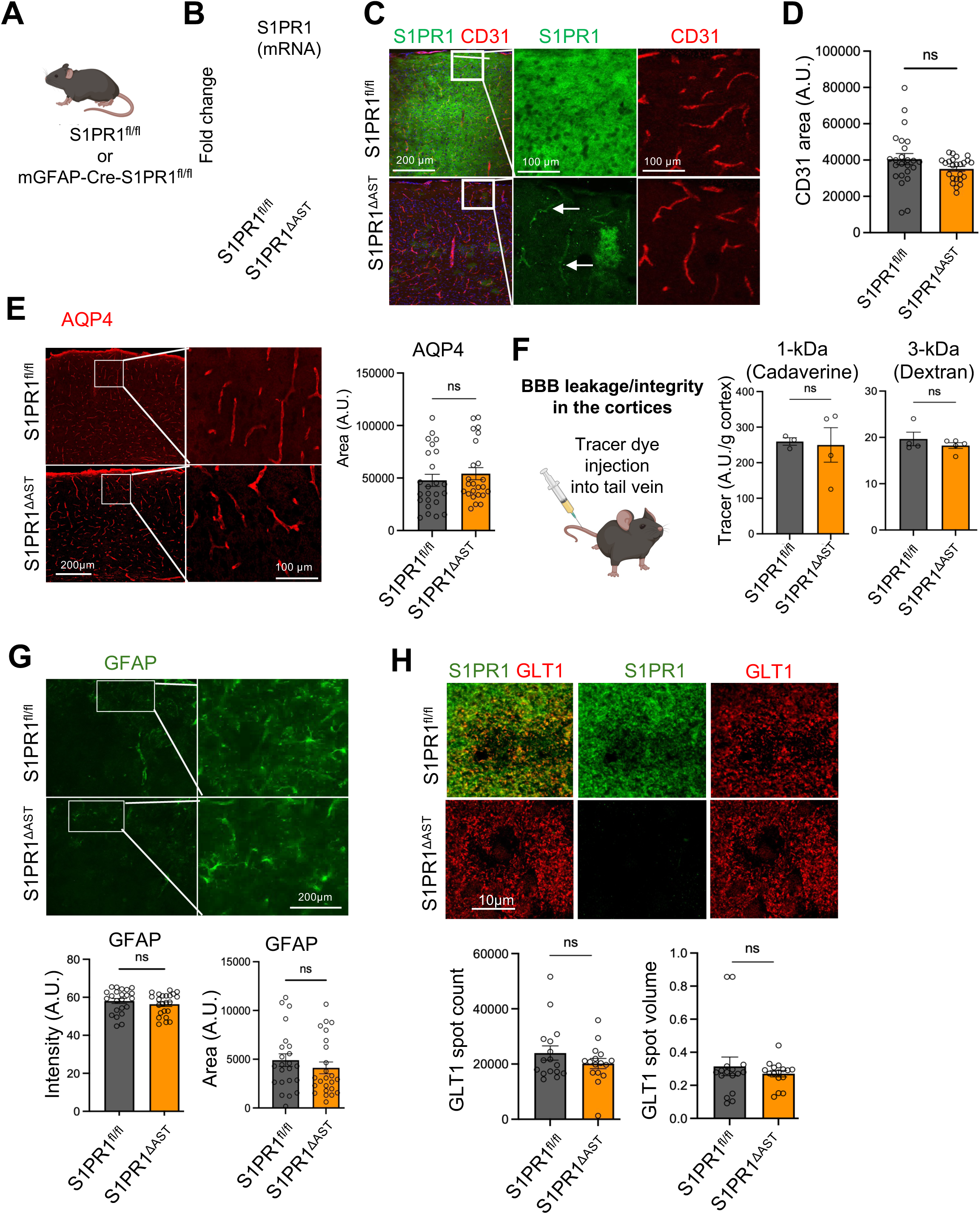
Astrocyte-specific, S1PR1 knock-out mice (S1PR1^ΔAst^) exhibit no gross changes in blood vasculature and astrocyte maturation. (**A**) Astrocyte-specific, S1PR1 knock-out mouse model (S1PR1^ΔAST^) model. (**B**) A substantial reduction in S1PR1 mRNA levels measured by qPCR. ((n=8 and 7 mice per group). **(C)** Representative confocal images of S1PR1 (green) with the endothelial marker CD31 (red) from S1PR1^ΔAST^ or littermate control cortices at P30. Arrows indicate intact S1PR1 expression on CD31+ vasculature. (**D**) Quantification of CD31 intensity represented as area per image is not altered in S1PR1^ΔAST^ cortices at P30. (**F**) Blood-brain barrier integrity assay scheme (left) and quantification of tracer dyes 1 kDa Cadaverine (center) and 3 kDa Dextran (right) signal in S1PR1^ΔAST^ cortices at ∼P35 indicates intact BBB integrity in S1PR1^ΔAST^ mice. (**G**) Representative image (top) and quantifications of GFAP signal (green) in S1PR1^ΔAST^ cortices at P30. (**H**) Representative image of perisynaptic astrocyte marker GLT-1 (red) with S1PR1 (green) in S1PR1^ΔAST^ cortices at P30 (top). Quantification of GLT1 puncta measured as GLT1-spot count and -spot volume by IMARIS 3D rendering indicate no changes in GLT1 expression in S1PR1^ΔAST^ cortices at P30. Data represents the mean ± SEM from n=3 and 4 mice per group (D-H). *** = *p* < 0.001, ns=not significant; Unpaired Two-tailed student’s T-test.

Since S1P signaling modulates cell growth and proliferation and that S1PR1 is expressed during astrocyte development, we measured astrocyte-cell numbers in the cortices of P30 S1PR1^ΔAST^ by immunostaining for SOX9 and OLIG2 and counting SOX9+OLIG2- nuclei as astrocytes as previously reported^38^(Fig. S2). No apparent difference in the number of SOX9+OLIG2- cells were observed between control and KO, indicating that astrocyte number was unaltered (Fig. S2B). Additionally, we measured the maturation and reactivity status of astrocytes in S1PR1^ΔAST^ mice by immunostaining for GFAP and observed no change in either fluorescent intensity or GFAP structure in the cortex (Fig. 3G). We have previously demonstrated that S1PR1 is highly expressed on perisynaptic astrocyte processes (PAPs) and strongly colocalizes with PAP marker, GLT1 (SLC1A2; glutamate transporter encoded by the gene *Slc1a2*), in the developing brain^28^. Therefore, we immunostained, imaged and quantified GLT1 puncta (or “spot”) count and spot volume using IMARIS 3D rendering to generate GLT1 spots from S1PR1^ΔAST^ brains (Fig. 3H). We observed no significant difference in the number or size of GLT1 spots in astrocytes between S1PR1^ΔAST^ or littermate controls (Fig 3H), suggesting that the loss of astrocytic S1PR1 does not alter GLT1 expression or GLT1 containing PAPs.

### S1PR1 KO astrocytes exhibit altered morphology and complexity *in vivo*

Because S1PR1 regulates neuronal contact-induced astrocyte morphogenesis *in vitro* (Fig. 1)^28^, we asked whether astrocyte morphogenesis is also impacted *in* vivo during postnatal development after loss of astrocytic S1PR1. We used an adeno-associated virus (AAV)-based approach to sparsely label and cytosolically fill cortical astrocytes with YFP in S1PR1^ΔAST^ mice (Fig. 4A). Briefly, P2-P3 S1PR1^ΔAST^ or littermate control pups were intracranially injected directly into the cortex with the AAV, pZac2.1-GfaABC_1_D-YFP, and brains were collected at P30, a timepoint where most astrocyte proliferation and maturation have completed in mice (Fig. 4A). Since astrocytes exhibit highly branched fine processes, 50 μm confocal Z-stacks of individual astrocytes were imaged and 3D reconstructed using IMARIS to generate surface and filament renderings for analysis (Fig. 4B).

**Fig 4.**
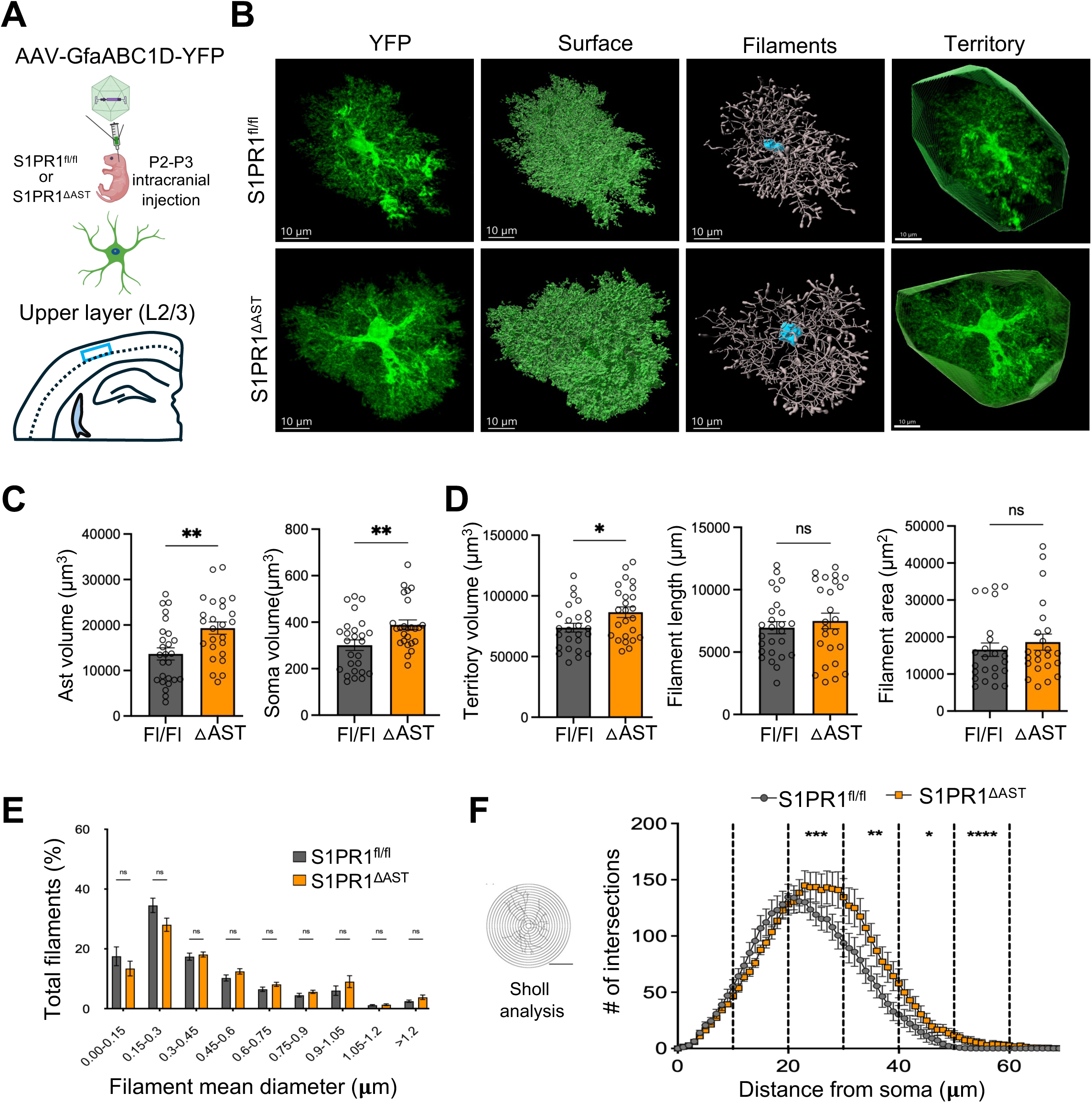
L2-3 S1PR1^ΔAST^ astrocytes exhibit increased morphology and complexity. (**A**) S1PR1^ΔAST^ or littermate controls were intracranially injected with the adenoassociated virus (AAV), pZac2.1-GfaABC1D-YFP, to sparsely label astrocytes. Sparsely labeled astrocytes from L2-3 somatosensory cortices were imaged and analyzed. (**B**) Representative confocal images and subsequent IMARIS 3D surface/filament renderings of YFP astrocytes from S1PR1^ΔAST^ or littermate controls at P30. **(C)** Quantification of astrocyte volume (left), and soma volume (right) from surface renderings generated using IMARIS. (**D**) Quantification of astrocyte territory volume (left), astrocyte process/filament length (middle) and area (right) from filament traces generated using IMARIS. (**E**) Plot depicting the proportion of astrocyte filament thicknesses grouped by mean diameter and represented as % of total filaments. (**F**) Sholl analyses of IMARIS rendered filament traces of astrocytes from S1PR1^ΔAST^ and littermate controls. Data represents the mean ± SEM from total 26 and 25 astrocytes from n=5 mice per group. * = *p*<0.05, *** = *p*<0.001, ns=non-significant, Unpaired Welch’s T-test.

Astrocytes are both functionally and structurally heterogeneous depending on brain region, neural input, and cell-cell interactions^4,11^. The cortex can be subdivided into different functional layers based on synaptic inputs and cytoarchitecture and contains diverse subpopulations of astrocytes that have been extensively characterized^36^. Therefore, we decided to focus our studies on the upper (L2-3) and deeper (L4-5) layer cortical astrocytes *in vivo*. Moreover, we chose the somatosensory cortex (SSC) for our analyses because a) SSC astrocytes are shown to modulate peripheral neuropathic pain and astrocytic S1PR1 is also crucial for neuropathic pain^37–39^ and b) to avoid potential injury-induced effects on astrocytes at the injection site in the visual cortex. In the absence of S1PR1, L2-3 astrocytes had larger territories as defined by their total volume, soma volume and territory volume (Fig. 4C,D), suggesting that S1PR1 limits astrocyte territory formation. Next, we performed filament tracing analyses on these astrocytes to precisely determine various parameters of their processes (branches) (Fig. 4D). S1PR1^ΔAST^ astrocytes showed no changes in total filament length nor total filament area (Fig. 4D). Further, analyses of astrocyte processes/filaments based on their thickness and grouped as a percentage of the total filament distribution (Fig. 4F) did not reveal any differences regarding the proportion of thicker (likely main processes) to thinner (distal processes or leaflets) processes per astrocyte (Fig. 4E). Although these results clearly demonstrate overall larger astrocyte territory without changes in processes parameters in S1PR1^ΔAST^, these analyses still do not show morphological differences in relation to the astrocyte soma. Therefore, we utilized three-dimensional Sholl analysis on the filament tracings to precisely plot astrocytic branch points (or intersections) in relation to the center of the astrocyte soma (Fig. 4F). S1PR1^ΔAST^ astrocytes displayed significantly larger intersections beyond 20 μm from the center, indicating increased branching in distal processes (Fig. 4F, right panel). Taken together, these results demonstrated that upper-layer cortical astrocytes are significantly larger in S1PR1^ΔAST^. Additionally, larger astrocytes are not due to a decrease in total number of astrocytes in S1PR1^ΔAST^ mice since quantification of astrocyte numbers (Sox9+Olig2-) in the cortical layers revealed no apparent change (Fig. S2).

Next, we performed identical analyses from deeper cortical layer (layers 4-5) astrocytes from S1PR1^ΔAST^ at P30 (Fig. 5A,B). Unlike astrocytes within upper layers of the cortex, deletion of S1PR1 had no major effect on L4-5 astrocyte size or territory (Fig. 5C, 5D). The total astrocyte volume, soma volume and territory volumes remained unchanged between cohorts (Fig. 5C, 5D) as did the average total filament area and filament length (Fig. 5D); however, Sholl analyses revealed a unique reduction in process branching between 20-40 μm away from the center of the cell body (Fig. 5F, left panel) while no differences were observed in the processes near the cell body and fine distal processes beyond 40 μm away, unlike in L2-3 astrocytes (Fig. 4F). Furthermore, these L4-5 astrocytes did not show any change in the process size distribution (Fig. 5E). Taken together, we can conclude that S1PR1^ΔAST^ possesses layer-specific morphological alterations.

**Fig 5.**
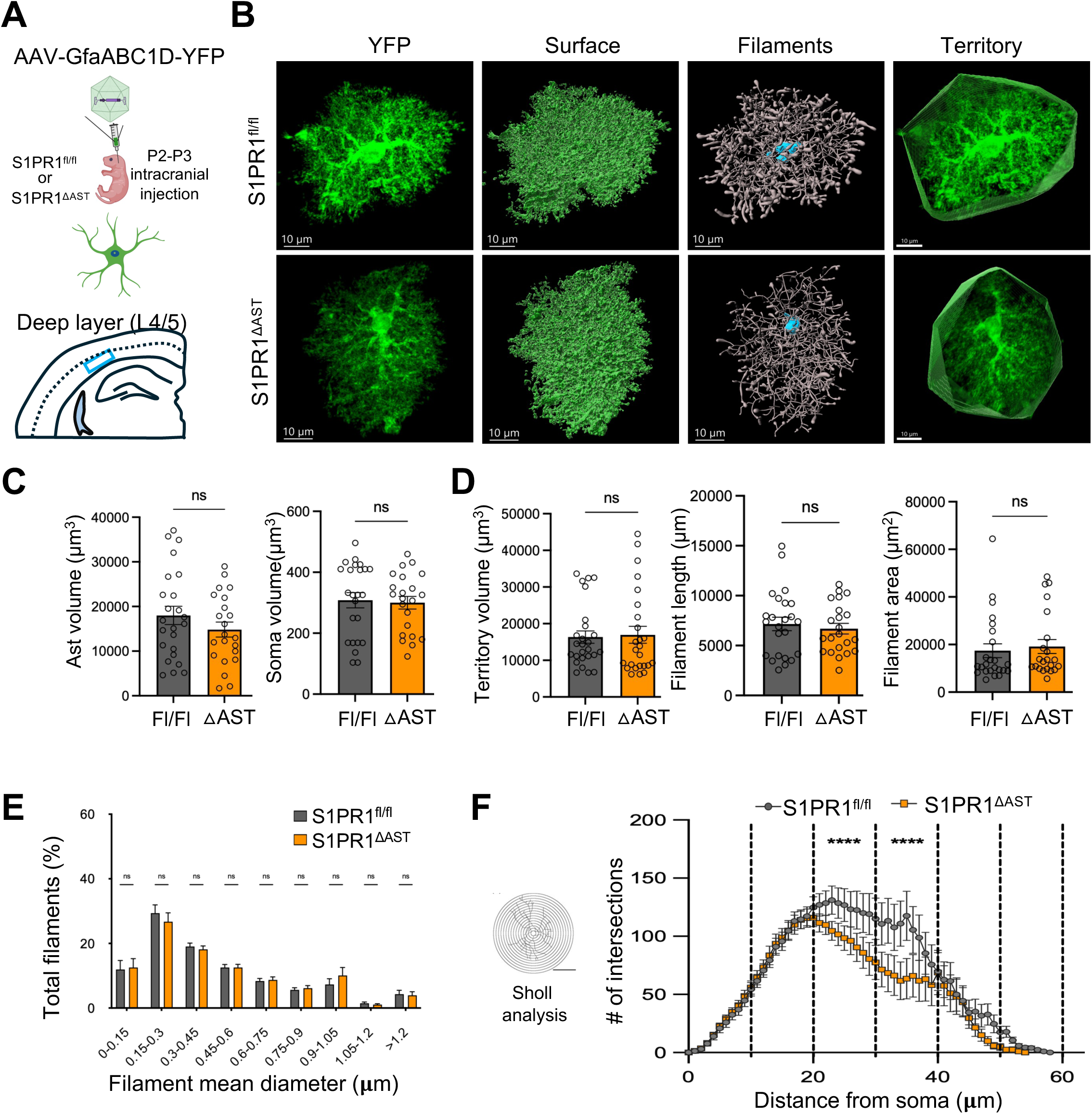
L4-5 S1PR1^ΔAST^ astrocytes exhibit a milder perturbation in astrocyte morphology. (**A**) Sparsely YFP-labeled L4-5 somatosensory cortical astrocytes were imaged and analyzed. **(B)** Representative confocal images and subsequent IMARIS 3D surface/filament renderings of YFP astrocytes from S1PR1^ΔAST^ or littermate controls at P30. **(C)** Quantification of astrocyte volume (left) and soma volume (right) from surface renderings generated using IMARIS. (**D**) Quantification of astrocyte territory volume (left), astrocyte process/filament length (middle) and area (right) from filament traces generated using IMARIS. (**E**) Plot depicting the proportion of astrocyte filaments grouped at various mean diameter and represented as % of total filaments revealed no differences in mean dendrites in S1PR1^ΔAST^ astrocytes. (**F**) Sholl analyses of IMARIS rendered filament traces of astrocytes from S1PR1^ΔAST^ and littermate controls. Data represents the mean ± SEM of 24 and 22 astrocytes from n=5 mice per group. * = *p*<0.05, ** = *p*<0.01, ns=non-significant, Unpaired Welch’s T-test.

### Membrane-tagged GFP filled S1PR1 KO astrocytes exhibit altered complexity *in vivo*

Astrocyte processes ramify into branches, branchlets, and leaflets, which contribute to 90–95% of an astrocyte’s total surface area and volume^11,40^. We reasoned that cytosolic GFP-labeling may only define larger astrocyte processes and may not fully encapsulate the morphology or territory occupied by a single astrocyte and its smaller/finer processes, and thus, may not detect effects of S1PR1 in these finer processes. Therefore, we labeled astrocyte membranes via an AAV mediated expression of Lck-GFP (pZac2.1 GfaABC_1_D-Lck-GFP), a membrane-tagged GFP, in S1PR1^ΔAST^ brains (Fig. S4) and performed the same analyses as in figures 4 and 5. Indeed, the volume, surface area, and number of intersections in Lck-GFP astrocytes were approximately 2-fold higher than that of cytosolic GFP filled astrocytes (compare Fig 4C and S4C) and are comparable to previously reported values for cortical astrocytes^36^. Lck-GFP-labeled S1PR1^ΔAST^ astrocytes showed no significant changes in volumes and surface area in both the upper and deeper cortical layers (Fig. S4 C,D and H,I). Sholl analyses of L2-3 Lck-GFP S1PR1^ΔAST^ astrocytes showed an increase in the number of intersections at distances greater than 20 microns from the cell body (Fig. S4E) while L4-5 Lck-GFP S1PR1^ΔAST^ astrocytes showed a significant decrease in the number of intersections between 15-30 microns from the cell body (Fig. S4J) The Sholl analyses are consistent between cytosolic YFP filled and membrane-tagged Lck-GFP experiments. Taken together these results suggest an increased complexity in L2-3 and a decreased complexity in L4-5 cortical astrocytes in S1PR1^ΔAST^ and overall reflect our initial observations in cytosolic YFP-labeled astrocytes in earlier analyses (Fig. 4,5), further solidifying a layer-specific difference in S1PR1-dependent astrocyte morphogenesis.

### Sparse knock-out of S1PR1 in the upper cortical layers reduces astrocyte morphology and complexity

Global loss of astrocytic S1PR1 *in vivo* led to an increase in astrocyte morphological complexity in the superficial layers (Fig. 4 and S4E), while there was a minor reduction in the deeper layers (Fig. 5, S4J). Our findings are in direct opposition to our findings *in vitro* (Fig. 1) and to those of others in the field studying invertebrate and zebrafish models^29^. We hypothesized that our findings here may represent a unique complexity in mammalian models of astrocyte morphology and, more specifically, astrocyte tiling. Although astrocyte tiling is conserved in mammals, zebrafish, and Drosophila, the molecular mechanisms driving this phenomenon are less known. It is, however, understood that astrocyte domains are established by repulsive or competitive interactions between neighboring astrocytes^11^. Therefore, we hypothesized that conditional loss of S1PR1 in the majority of astrocytes would lead to loss of competition-driven tiling and extended astrocyte processes and territory volume. We predicted that a difference in S1PR1 expression in neighboring astrocytes would result in reduced morphological complexity and territory in the astrocytes with reduced S1PR1 expression (Fig. 6B). To test this possibility, we simultaneously fluorescently labeled and sparsely deleted S1PR1 in cortical astrocytes by injecting either the astrocyte-specific AAV pZac2.1 GfaABC1D-YFP-P2A-Cre or YFP as a control into S1PR1^loxP/loxP^ pups at P2-3 (Fig. 6A-C). We validated sparse S1PR1 knockout in YFP-P2A-Cre expressing astrocytes at P30 and performed the exact same imaging parameters and morphometric analyses as in Fig. 4,5.

**Fig 6.**
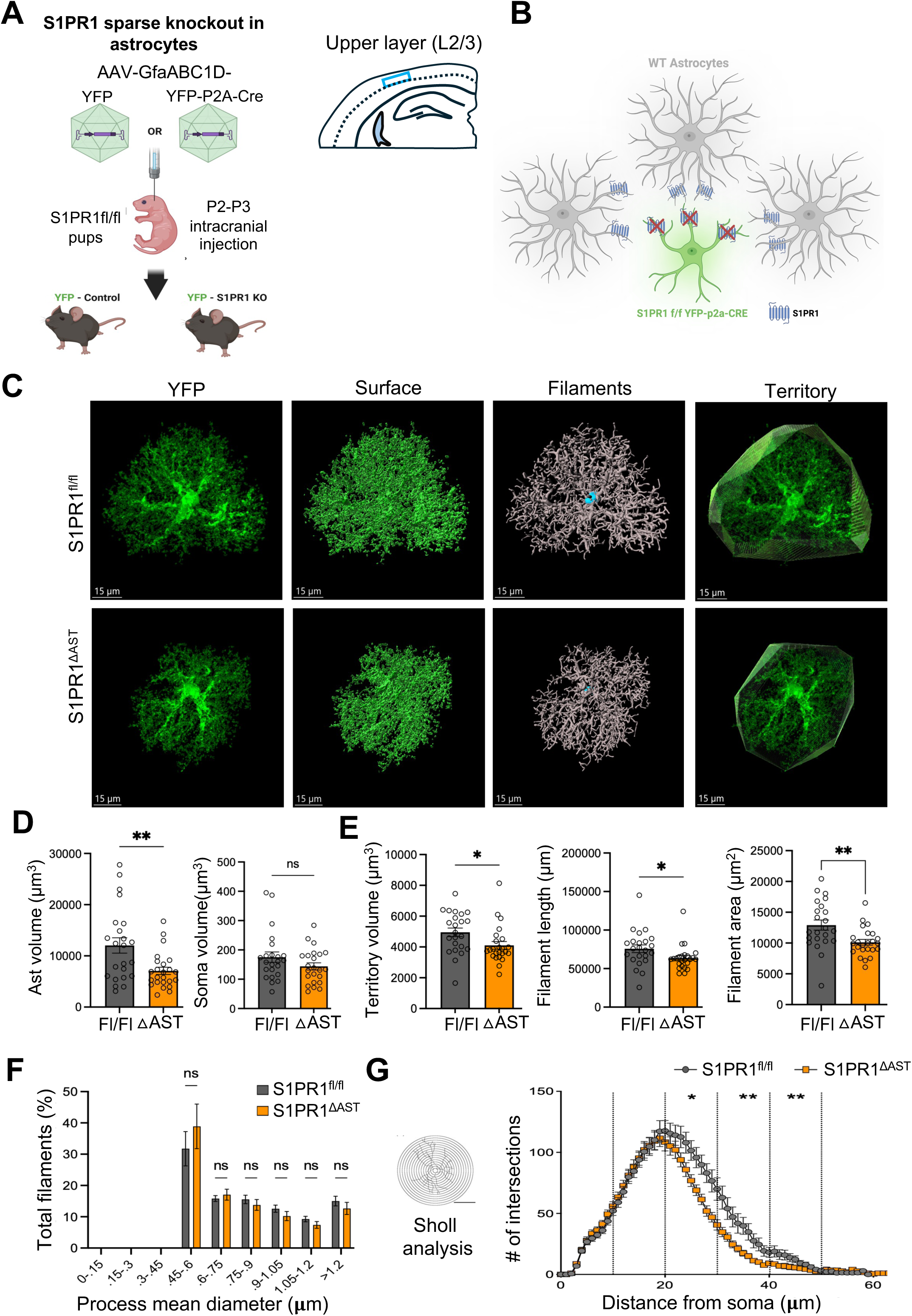
Sparse deletion of S1PR1 in L2-3 astrocytes results in reduced astrocyte size and morphological complexity. **(A)** Schematics showing S1PR1 deletion in sparse astrocytes by delivering AAV-GfaABC1D-YFP-P2A-Cre or AAV-GfaABC1D-YFP control in S1PR1^fl/fl^ mouse pups. **(B)** Sparsely labeled astrocytes were imaged from L2-3 somatosensory cortex from P30 pups. (**C**) Representative confocal images and subsequent IMARIS 3D surface/filament renderings of YFP astrocytes from S1PR1^ΔAST^ or littermate controls at P30. **(D)** Quantification of astrocyte volume (left) and soma volume (right) from Surface renderings generated by IMARIS. (**E**) Quantification of astrocyte territory volume (left) filament length (middle) and filament area (right) from filament traces generated by IMARIS. **(F)** Plot depicting the proportion of astrocyte filaments grouped by various mean diameter and represented as % of total filaments revealed no differences in mean dendrites in S1PR1^ΔAST^ astrocytes. (**G**) Sholl analyses of IMARIS rendered filament traces of YFP labeled sparse astrocytes from S1PR1^ΔAST^ and littermate controls. Data represents the mean ± SEM of 23 and 24 astrocytes from n=4 mice per group. * = *p*<0.05, ** = *p*<0.01, ns=non-significant, Unpaired Welch’s T-test.

Interestingly, individual S1PR1 KO (YFP-P2A-Cre injected) astrocytes in the upper layers exhibited reduced astrocyte volume (Fig. 6D), territory volume (Fig. 6E), and a reduction in complexity in the finer processes measured as number of intersections by Sholl (Fig. 6G) compared to control astrocytes (YFP injected only). Additionally, we saw a significant reduction in filament length and area (Fig. 6E) but no obvious change in filament diameter populations (Figs. 6F). Collectively, these results indicate that sparse S1PR1 KO astrocytes are overall smaller and less complex in the upper cortical layers. These results contradict our previous morphometric results from S1PR1 deletion in a total astrocyte background (Figs. 4) but are in line with our prediction that S1PR1 may play a role in competition-dependent astrocyte growth and tiling of the upper cortical layers (Fig. 6B).

Next, we focused our analyses on deeper layer astrocytes in the sparse S1PR1 KO model (Fig. 7A,B). Deletion of S1PR1 from individual astrocytes in the deeper layer showed a strong reduction in overall astrocyte volume and territory volume compared to control astrocytes (Fig. 7C,D). Moreover, Sparse S1PR1 deletion from deeper layer astrocytes also exhibited a reduction in the number of intersections compared to control (Fig. 7F); however, no significant redistribution in process diameters or overall process length was observed (Fig. 7E). Since excessive Cre expression may affect cellular health and affect astrocyte morphology even in the absence of a floxed allele, we injected AAVs expressing YFP or YFP-P2A-Cre into C57BL/6J (wild-type) pups at P2-3 and subsequently analyzed astrocyte morphology (Fig. S5). In our hand, Cre expression alone did not cause any substantial changes in astrocyte volume, territory or branching (Fig. S5C-E). Taken together, sparse S1PR1 KO astrocytes in deeper cortical layers showed an overall reduced territory size and stronger reduction in complexity, suggesting a role of S1PR1 in competition-driven astrocyte process growth and tiling.

**Fig 7.**
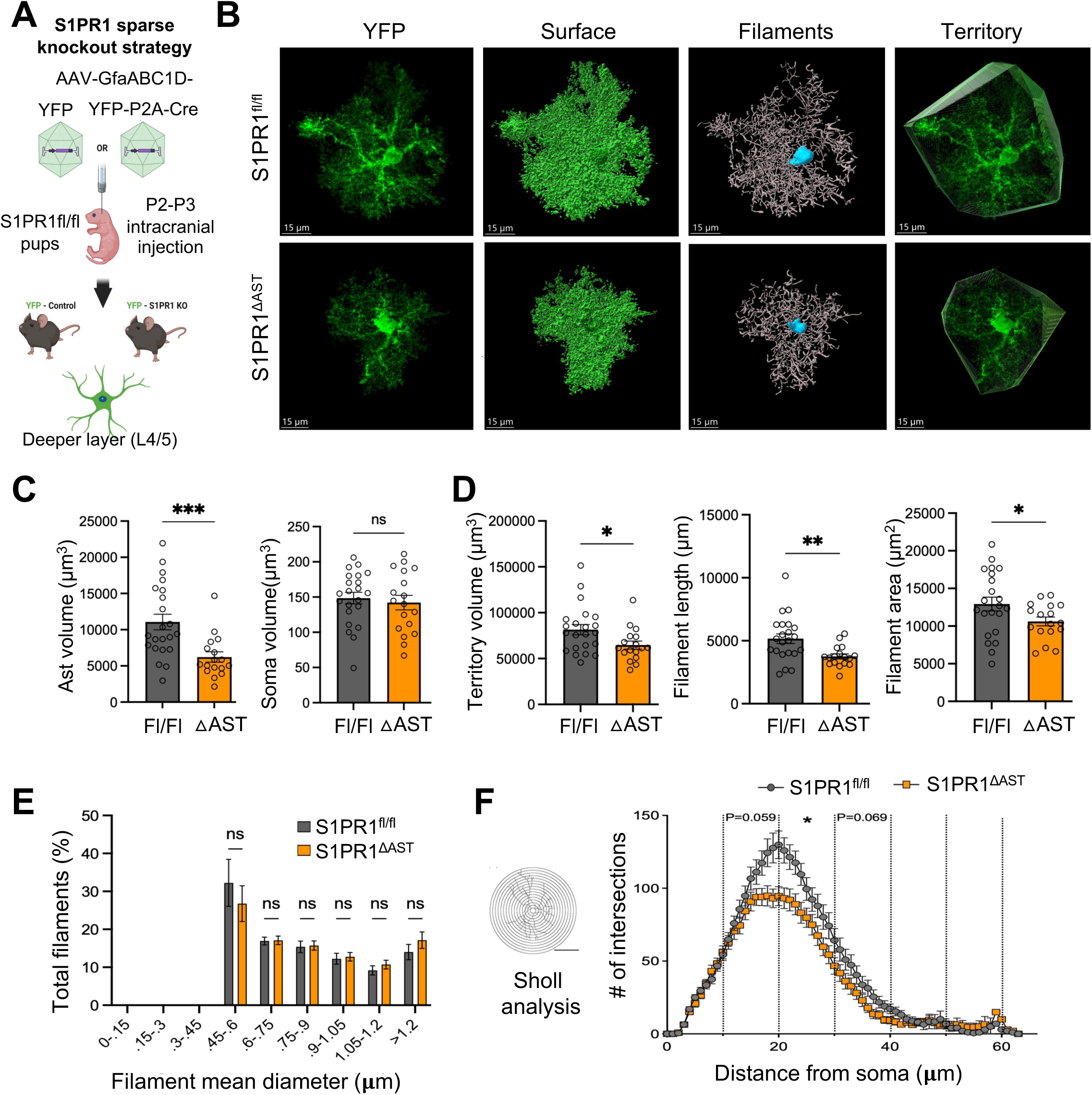
Sparse deletion of S1PR1 in L4-5 astrocytes results in reduced astrocyte size and morphological complexity. **(A)** Schematics showing S1PR1 deletion in sparse astrocytes by delivering AAV-GfaABC1D-YFP-P2A-Cre or AAV-GfaABC1D-YFP control in S1PR1^fl/fl^ mouse pups. Sparsely labeled astrocytes were imaged from L4-5 somatosensory cortex from P30 pups. (**B**) Representative confocal images and subsequent IMARIS 3D surface/filament renderings of YFP astrocytes from S1PR1^ΔAST^ or littermate controls at P30. **(C)** Quantification of astrocyte volume (left) and soma volume (right) from Surface renderings generated by IMARIS. (**E**) Quantification of astrocyte territory volume (left) process/filament length (middle) and area (left) from filament traces generated by IMARIS. **(F)** Plot depicting the proportion of astrocyte filaments grouped by mean diameter and represented as % of total filaments revealed no differences in mean dendrites in S1PR1^ΔAST^ astrocytes. (**G**) Sholl analyses of IMARIS rendered filament traces of GFP labeled sparse astrocytes from S1PR1^ΔAST^ and littermate controls. Data represents the mean ± SEM of 22 and 17 astrocytes from n=4 mice per group. * = *p*<0.05, ** = *p*<0.01, *** = *p*<0.001 ns=non-significant, Unpaired Welch’s T-test.

### S1PR1 regulates astrocyte territorial overlap/tiling

So far, our results indicate a role for astrocytic S1PR1 in competition-dependent astrocyte tiling. To specifically test this, we deleted S1PR1 in two adjacent astrocytes in cortical areas by coinjecting AAVs expressing YFP-P2A-Cre and tdTomato-P2A-Cre (KO:KO) or YFP and tdTomato (WT:WT) under the control of the GfaABC_1_D promoter into S1PR1^loxP/loxP^ pups at P2-3 (Fig. 8A-C). Although this strategy creates coinfection of YFP and tdTomato mostly within the same astrocyte, in some instances an astrocyte is infected/labeled with only one (YFP or tdTomato) fluorescent protein (Fig. 8B-D). Furthermore, in rare instances we found two adjacent astrocytes labeled with YFP and tdTomato ideal for analyzing territorial overlap/tiling (we called these as “Tiling pair” astrocytes) (Fig. 8D). After searching through several brain slices from 3 mice per group we found and imaged ∼18-23 tiling pair astrocytes. While we found minimal territorial overlap amongst the WT:WT tiling pairs, astrocytes in the S1PR1 KO:KO tiling pair showed extensive territorial overlap volume relative to their total volume captured (Fig. 8D,E). This data clearly demonstrates that S1PR1 regulates astrocyte territorial volume overlap, causing a tiling defect in a competition-dependent manner.

**Fig 8.**
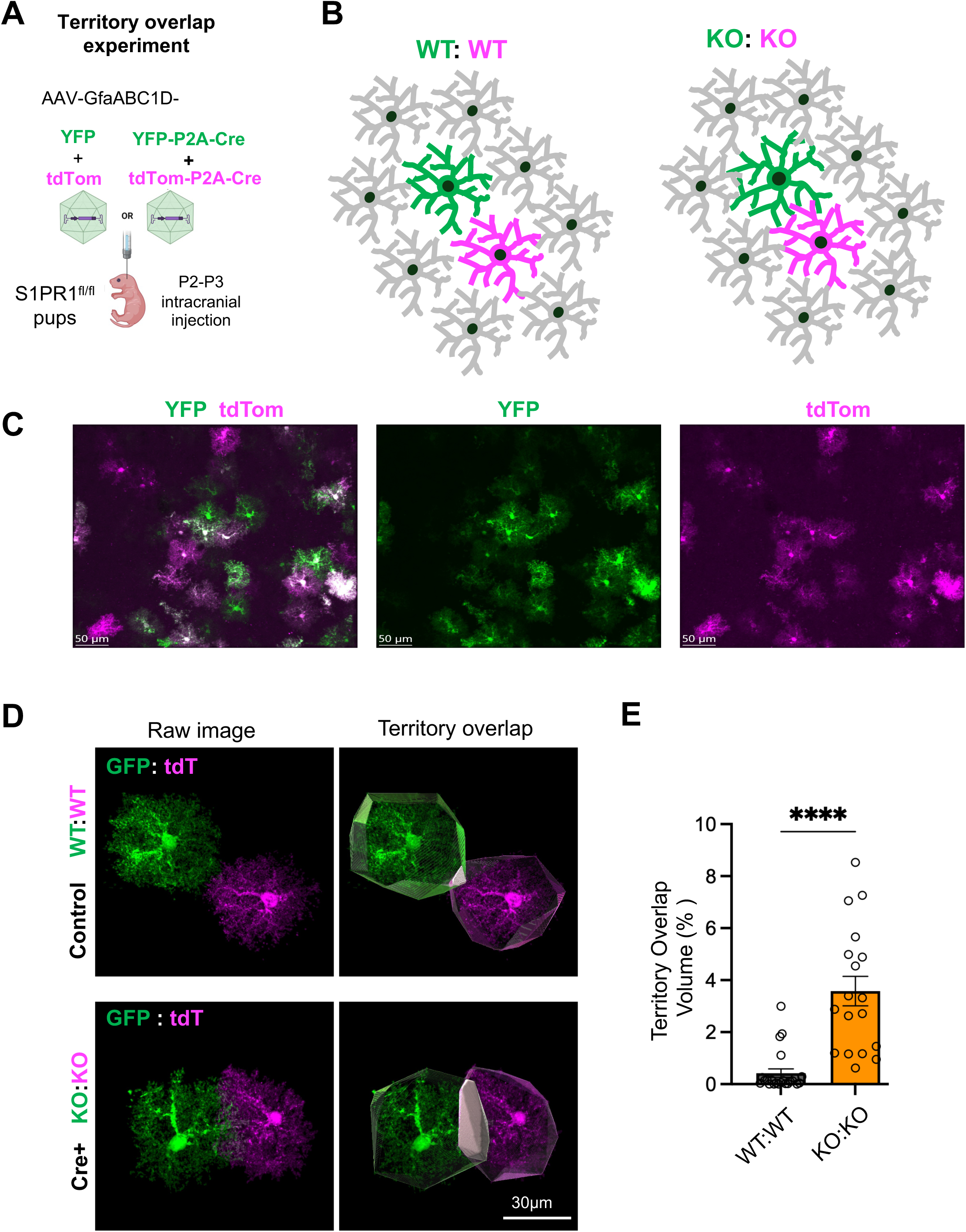
Neighboring S1PR1 KO astrocytes exhibit increased astrocyte territory overlap. (A, **B)** Schematics showing S1PR1 deletion in neighboring astrocytes in two-color schemes by delivering AAV-GfaABC1D-YFP-P2A-Cre plus AAV-GfaABC1D-tdTomato-P2A-Cre or AAV-GfaABC1D-YFP plus AAV-GfaABC1D-tdTomato controls in S1PR1^fl/fl^ mouse pups. (**C**) Example 20x confocal image to show the labelling by two color viruses. **(D)** Representative confocal images and subsequent IMARIS 3D surface/filament renderings to create territories of neighboring astrocytes from KO:KO or littermate WT:WT controls at P30. **(E)** Quantification of astrocyte territory overlap volume represented as % of total volume from two neighboring astrocytes. Data represents the mean ± SEM of 23 and 18 pairs of neighboring astrocytes from n=3 and 4 mice per group. **** = *p*<0.0001, Unpaired Welch’s T-test.

## Discussion

Advancements in molecular profiling and super-resolution microscopy have identified astrocytes as heterogenous cells that possess unique molecular, transcriptional, and morphological characteristics^41,42^. Their tumultuous morphology enables them to engage in numerous neurological functions through interactions with neurons, glia, and vasculature at both circuit and synaptic levels. Thus, uncovering the mechanisms that establish these interactions is essential to understanding the development and function of the mammalian brain. How astrocyte morphogenesis and tiling are established in the mammalian brain is mostly unknown. Here we demonstrate that the astrocyte-enriched G-protein coupled receptor S1PR1 influences morphogenesis of murine astrocytes both *in vitro* and *in vivo*. Moreover, our data strongly suggest that S1PR1 may play a role in establishing competition-driven astrocyte morphogenesis and tiling and further propose that the bioactive lipid S1P drives astrocyte growth and tiling in a neuronal contact-dependent manner.

### Neuronal contact induces S1PR1-dependent astrocyte morphogenesis *in vitro*

In mouse astrocyte-neuron cocultures, we observed two distinct pools of astrocytes. One with high S1PR1 and more complex morphology (astrocytes in contact with neurons) and the other with low S1PR1 and less complex morphology (astrocytes without contact with neurons). This strongly suggests that a contact-dependent signal, a labile soluble signal, or an extracellular matrix component released or deposited locally by neurons may induce S1PR1 expression and astrocyte morphogenesis. Indeed, our inhibitor data demonstrate that JAK-STAT3 but not SHH signaling regulates S1PR1 expression and morphological complexity (Fig. 2). Although SHH is a soluble signal released by deep-layer cortical neurons, it interacts with and activates PTCH1 in astrocytes localized near the SHH expressing neuronal cell body^33^. Similarly, CT1 could also be locally released by neurons to induce JAK-STAT3 signaling in astrocytes to regulate S1PR1 expression and astrocyte morphogenesis. Because neurexin-neuroligin or delta-catenin-based cell adhesion interactions between neuron and astrocyte have been shown to regulate astrocyte morphogenesis, their involvement in S1PR1 expression cannot be ruled out here^30,43^. Interestingly, S1P-S1PR1 axis has been shown to induce JAK-STAT3 signaling in cancer cells and that S1PR1 antagonist W146 inhibited expression of S1PR1 in cocultured astrocytes (Fig. 1K), it is possible that neuronal-derived S1P could induce S1PR1 expression in cocultured astrocytes by enhancing JAK-STAT3 signaling. Because S1P can induce SHH expression through JAK–STAT3 activation in other cell types^46^, it is plausible that neuron-derived S1P similarly activates JAK–STAT3 in neurons to promote SHH production, which then drives contact-dependent astrocyte morphogenesis. In parallel, S1P directly signals through S1PR1–JAK–STAT3 in astrocytes to enhance their morphological complexity. Thus, two distinct S1P-dependent pathways—one neuronal (S1P→JAK–STAT3→SHH) and one astrocytic (S1P→S1PR1→JAK–STAT3)—may jointly shape astrocyte development in our cocultures. This dual-pathway model also explains why Ruxolitinib, which inhibits JAK–STAT3 in both neurons and astrocytes, produces a stronger reduction in astrocyte complexity and S1PR1 expression than Sonidegib, which disrupts only the SHH-dependent astrocyte complexity.

### S1PR1 regulates astrocyte morphogenesis in a cortical-layer dependent manner

AAV-mediated tracing of cytoplasm-filled or membrane-tagged GFP filled cortical astrocytes revealed layer-specific differences in S1PR1-dependent astrocyte morphogenesis. We observed that S1PR1 expression influenced the size, territory, and complexity of astrocytes in upper cortical layers, whereas astrocytes in deeper layers appeared modestly affected. In agreement with our findings, recent work has identified that cortical astrocytes are molecularly, structurally, and functionally heterogeneous in a layer-specific manner^36,44^. Formation of the cortex occurs in a well-described “inside-first, outside-last” pattern, where neural progenitor cells migrate to, proliferate, and differentiate into neurons in deeper layers prior to upper layers^45,46^. Mouse astrocytes begin to differentiate and interact with neurons toward the conclusion of embryogenesis and the onset of corticogenesis; however, they do not achieve structural or functional maturation until 3–4 weeks post-birth^5^. Notably, the expression of S1PR1 coincides with corticogenesis and astrocyte maturation, gradually increasing during postnatal development and plateauing around P30^28^. Given the temporal overlap between corticogenesis and S1PR1 expression, it is possible that deeper layer astrocytes, which have formed connections with neurons, have already structurally matured at P30 and no longer depend on S1PR1 activity. If true, deeper layer astrocytes would exhibit morphological deficits at earlier developmental stages in the absence of S1PR1.

Alternatively, layer-specific differences in S1PR1-dependent astrocyte morphogenesis may stem from inherent variations in basal S1PR1 expression across cortical layers. Spatial transcriptomic analyses of the mouse cortex have revealed that S1PR1 is enriched in a subpopulation of astrocytes localized within upper cortical layers^47^. Our immunofluorescent staining shows that S1PR1 appears to have the strongest signal in upper cortical layers when compared to deeper layers (Fig. 3C and data not shown). Consequently, higher expression of S1PR1 in upper-layer astrocytes might correlate with increased S1PR1 activity. This suggests that the S1P-S1PR1 signaling axis could be more pronounced within upper cortical layers, which could explain our layer-specific differences.

Astrocyte morphogenesis is governed by a range of redundant molecular mechanisms that vary by cortical layer. Cell-cell adhesion, neural activity, and neuron-secreted growth factors have been described in regulating astrocyte morphogenesis and have been shown to be relevant in different cortical layers^43,48,49^. For instance, loss of the signaling molecule Sonic hedgehog (Shh) in layer-5 neurons reduced astrocyte complexity and tripartite synapse coverage; conversely, cell-autonomous activation of Shh signaling in astrocytes promoted cortical excitatory synapse formation^33^. Additionally, catenin-cadherin-mediated cell adhesion between neurons and astrocytes in deeper cortical layers, not upper, established astrocyte morphological complexity^43^. Together, these findings underscore the existence of compensatory, layer-specific, and bidirectional communication mechanisms that may mitigate the impact of S1PR1 loss in deeper layer astrocytes.

### S1PR1 and Astrocyte Tiling

Contrary to S1PR1^ΔAST^ astrocytes, deletion of S1PR1 from individual or sparse populations of cortical astrocytes caused a reduction in astrocyte size, territory, and complexity, regardless of cortical layer (Fig 6,7). Although consistent with our *in vitro* findings, these results contradict our initial *in vivo* observations and raise the question: why does the ablation of S1PR1 in select astrocytes result in a decrease in their overall size? A potential explanation could be competition. Astrocytes occupy distinct, non-overlapping territories with neighboring astrocytes. Within a single astrocyte territory, astrocytes have at least one main process that associates with blood vessels and hundreds of finer processes that can contact thousands of synapses^50,51^. These domains are formed by competitive, activity-dependent, and repulsive mechanisms that limit an astrocyte’s size, shape, and overall complexity^11,52^—a phenomenon referred to as tiling. Growth factor and metabolite concentration gradients common in the brain^53^ are used to drive chemotaxis of astrocyte processes and neurites during postnatal brain development^54,55^. In the immune system, maintenance of an S1P gradient between lymphatic tissue and the blood is important for the extravasation of immune cells in an S1PR1-dependent manner^56^. Extracellular S1P levels are regulated by the degradation of S1P into sphingosine via phospholipid phosphatases and subsequent internalization^57,58^. Interestingly, activity of the lipid phosphatases Wunen (wun) and Wunen2 (wun2) are implicated in S1PR1-dependent astrocytic tiling in zebrafish^29^. Furthermore, it has been postulated that S1PR1 recycling is necessary for buffering extracellular S1P^59^. This behavior is likely due to the docking of S1P’s lipophilic sphingoid base into the binding cavity of S1PR1 within the lipid membrane, followed by the subsequent endocytosis and degradation of S1P-S1PR complexes^60^. This evidence indicates that lipid phosphatases and S1P receptors can control levels of extracellular S1P and influence astrocyte process dynamics. Thus, in our sparse-S1PR1 knockout model, a lack of S1PR1 could influence astrocyte morphogenesis by 1) regulating cytoskeleton remodeling through S1PR1; and 2) indirectly promoting an excess of extracellular S1P, which could favor adjacent wild-type astrocyte process outgrowth. However, the intercellular mechanisms of S1P release, buffering, and degradation within the brain are still under investigation.

### Potential Mechanisms of S1PR1 in Astrocyte Morphogenesis

From lamellipodia formation to tumor cell metastasis, S1P-S1PR signaling has been implicated in a diverse array of biomechanical processes^61–64^. In the mouse brain, S1PR1 primarily localizes to fine astrocytic PAPs, which act as interfaces for astrocyte-neuron cross-communication^28^. The remodeling of these PAPs in response to neuronal activity fine-tunes the regulation of neurotransmitter clearance, ion exchange, and metabolite transport. S1PR1 has been directly associated with modulation of Rac and Rho GTPase activity, molecular switches that regulate actin filament dynamics necessary for astrocyte process motility^65,66^. Furthermore, deletion of S1PR1 or inactivation of sphingosine kinase 1 (SphK1)—one of the enzymes that converts sphingosine into S1P—suppresses Rac1 activity and limits chemotaxis^67,68^. Therefore, the localization of S1PR1 on PAPs positions the receptor as a potential regulator of PAP remodeling in response to environmental and molecular cues such as neuronal activity^69^. Indeed, S1PR1 has been shown to regulate Rac1-mediated actin remodeling of astrocyte processes in zebrafish, leading to dynamic membrane extension and retraction^29^, deficits in which generated smaller astrocytes. In support of this, our results indicate that S1PR1 influences murine astrocyte size, territory, and complexity, likely by regulation of process branching through actin remodeling. Moreover, S1PR1 KO astrocytes mostly exhibited changes in finer, distal processes regardless of the KO model, as evidenced by Sholl analyses (Figures 4-7 and S4). Thus, S1PR1 might be primed to regulate dynamic, actin-dependent cytoskeletal remodeling in finer astrocytic processes.

### Consequences and Future Directions

The formation and remodeling of fine astrocyte processes are essential for maintaining normal astrocyte function, aberrations in which precede many neurological diseases^70–72^. Astrocytes engage in dynamic interactions with both neurons and other glial cells to support the assembly, maintenance, and functionality of neural circuits. Our findings highlight a critical role for S1PR1 in the structural maturation of astrocytes; however, the underlying intercellular mechanisms driving this process remain poorly understood. Our in vitro experiments suggest that neuronal contact induces astrocytic S1PR1 expression, while exogenous S1P stimulation enhances astrocyte morphological complexity and upregulates synaptogenic factor expression^28^. These results raise the possibility that neuron derived S1P may act in a paracrine manner on astrocytic S1PR1 to promote tripartite synapse formation. Nonetheless, further investigation is needed to determine the cellular source of S1P *in vivo* and to elucidate the functional consequences of disrupted S1PR1 signaling during normal brain development and disease.

## Declarations

### Authors’ contributions

C.T., J.P.G. and S.C.R.T. planned and performed most experiments, with assistance from Z.T., S.M., P.M., J.H., K.S. and O.D. SKS conceived the study and contributed to planning of the experiments. C.T., J.P.G. and S.K.S. drafted the manuscript. All authors read and approved the final manuscript.

### Availability of data and material

The data that support the findings of this study are available from the corresponding author upon reasonable request.

### Ethics approval

Mice were housed at Virginia Commonwealth University according to guidelines of the Institutional Animal Care Use Committee (IACUC). The mouse protocols were approved by the IACUC. All mice were housed with food and water available ad libitum under a 12 h–12 h light–dark cycle in a 20–22°C and 40–60% humidity environment.

### Consent for publication

Not applicable.

## Supplementary figure legends

**Fig S1. Astrocyte, neuron isolation and coculture from S1PR1-GFP reporter mice and WT mice respectively to study expression and function of S1PR1. (A)** Schematics of isolation and culture of astrocyte and neurons in vitro. **(B)** Purity of astrocyte cultures and expression of S1PR1-GFP (green) appears plasma membrane and stained with astrocyte marker S100b (red) in astrocyte only and A+N cocultures. **(C)** S1PR1-GFP expressing astrocytes showed two population of cells bright with complex morphology vs dim and roundish less complex cellular morphology.

**Fig S2. Astrocyte numbers as quantified by SOX9+ and Olig2- remain unchanged in S1PR1^ΔAST^ mice. (A)** Representative confocal 20x images stained with SOX9 (red) OLIG2 (green) and Dapi (blue) from somatosensory cortices of **S1PR1^ΔAST^** or littermate control mice at P30. Boxed area is zoomed in and presented at the lower pannels. Arrowheads indicate SOX9+OLIG2- astrocytes. **(B)** Quantification of SOX9+OLIG2- astrocytes (left), SOX9+ cells (middle and OLIG2+ cells (right) represented as the cells/ 10,000μm^2^ showing no changes in astrocyte numbers. Data represents the mean ± SEM of 19 and 19 images n=3 and 3 mice per group. n.s=non-significant, Unpaired Welch’s T-test.

**Fig S3. GFAP-Cre dependent GFP reporter mouse display minimal reporter expression in non-astrocytic cells. (A)** Schematics to show our breeding strategy to generate Cre dependent reporter mouse line by crossing our GFAP-Cre (no flox) with Lox-STOP-Lox-Cas9-P2A-GFP mouse line. **(B)** Representative confocal 20x images stained with GFP (green) and costained with either neuronal marker NeuN(red) (arrow indicates a NeuN+ GFP+ neuron), or microglial marker Iba1 (red), or Oligodendrocyte lineage marker OLIG2 (red) or **(C)** astrocyte marker GFAP (red) from P24 cortices or Hippocampus (only GFAP/GFP **(D)). (E)** Table with quantification of GFP+NeuN+, or GFP+Iba1+ or GFP+OLIG2+ cells as % of GFP labeled cells. Data represents the mean ± SEM of 9-11 images from n=3 mice.

**Fig S4. Membrane-targeted Lck-GFP labeled astrocytes from S1PR1^ΔAST^ mice exhibit complexity changes.** (**A**) Schematics showing AAV-based membrane labeling of L2-3 astrocytes *in vivo* by Lck-GFP. **(A-F) Sparsely labeled astrocytes from L2-3 somatosensory cortices were imaged and analyzed.** (**B**) Representative, confocal L2-3 membrane targeted-GFP filled astrocytes (left panel) and IMARIS rendered surface traces (right panels) **(C)** IMARIS rendered filament traces and astrocyte territory. **(D)** Quantification of astrocyte volume from Surface renderings generated using IMARIS. (**E**) Quantification of astrocyte territory volume (left) and filament length from filament traces generated using IMARIS. (**F**) Sholl analyses of IMARIS rendered filament traces of Lck-GFP labeled astrocytes from S1PR1^ΔAST^ and littermate controls. Data represents the mean ± SEM of 25-28 astrocytes from n=3 mice per group. * = *p*<0.05, ** = *p*<0.01, ns=non-significant, Unpaired Welch’s T-test. (**G-L**) **Sparsely labeled astrocytes from L4-5 somatosensory cortices were imaged and analyzed.** (**H**) Representative, confocal L4-5 membrane targeted-GFP filled astrocytes (left panel) and IMARIS rendered surface traces (right panels) **(I)** IMARIS rendered filament traces and astrocyte territory. **(J)** Quantification of astrocyte volume from Surface renderings generated using IMARIS. (**K**) Quantification of astrocyte territory volume (left) and filament length from filament traces generated using IMARIS. (**L**) Sholl analyses of IMARIS rendered filament traces of Lck-GFP labeled astrocytes from S1PR1^ΔAST^ and littermate controls. Data represents the mean ± SEM of 27 and 37 astrocytes from n=3 mice per group. * = *p*<0.05, ** = *p*<0.01, **** = *p*<0.0001, ns=non-significant, Unpaired Welch’s T-test.

**Fig S5. Expression of Cre by AAV does not alter astrocytes morphology.** (**A**) Schematics showing delivery of AAV expressing GfaABC1D-YFP-P2A-Cre or GfaABC1D-YFP control in wildtype (WT) mouse pups. Sparsely labeled astrocytes were imaged from both upper and deeper somatosensory cortex from P30 pups. **(B)** Representative confocal images and subsequent IMARIS 3D surface/filament renderings of YFP astrocytes from control or Cre expressing littermates at P30. **(C)** Quantification of astrocyte volume (left) and soma volume (right) from Surface renderings generated using IMARIS. (**D**) Quantification of astrocyte territory volume from filament traces generated using IMARIS. (**E**) Sholl analyses of IMARIS rendered filament traces of astrocytes from Cre.P2A.YFP and YFP only. Data represents the mean ± SEM of 11 and 11 astrocytes from n=3 mice per group. ns=non-significant, Unpaired Welch’s T-test.

## References

(1) Zuchero, J. B.; Barres, B. A. Glia in Mammalian Development and Disease. Dev. Camb. 2015, 142 (22), 3805–3809. 10.1242/dev.129304.

(2) Xiong, H.; Tang, F.; Guo, Y.; Xu, R.; Lei, P. Neural Circuit Changes in Neurological Disorders: Evidence from in Vivo Two-Photon Imaging. Ageing Res. Rev. 2023, 87, 101933. 10.1016/j.arr.2023.101933.

(3) De Luca, C.; Colangelo, A. M.; Virtuoso, A.; Alberghina, L.; Papa, M. Neurons, Glia, Extracellular Matrix and Neurovascular Unit: A Systems Biology Approach to the Complexity of Synaptic Plasticity in Health and Disease. Int. J. Mol. Sci. 2020, 21 (4), 1539. 10.3390/ijms21041539.

(4) Degl’Innocenti, E.; Dell’Anno, M. T. Human and Mouse Cortical Astrocytes: A Comparative View from Development to Morphological and Functional Characterization. Front. Neuroanat. 2023, 17. 10.3389/fnana.2023.1130729.

(5) Akdemir, E. S.; Huang, A. Y.-S.; Deneen, B. Astrocytogenesis: Where, When, and How. F1000Research 2020, 9, F1000 Faculty Rev-233. 10.12688/f1000research.22405.1.

(6) Chaboub, L. S.; Deneen, B. Developmental Origins of Astrocyte Heterogeneity: The Final Frontier of CNS Development. Dev. Neurosci. 2013, 34 (5), 379–388. 10.1159/000343723.

(7) Bélanger, M.; Allaman, I.; Magistretti, P. J. Brain Energy Metabolism: Focus on Astrocyte-Neuron Metabolic Cooperation. Cell Metab. 2011, 14 (6), 724–738. 10.1016/j.cmet.2011.08.016.

(8) Sun, M.; You, H.; Hu, X.; Luo, Y.; Zhang, Z.; Song, Y.; An, J.; Lu, H. Microglia–Astrocyte Interaction in Neural Development and Neural Pathogenesis. Cells 2023, 12 (15), 1942. 10.3390/cells12151942.

(9) Lee, S. Y.; Chung, W.-S. The Roles of Astrocytic Phagocytosis in Maintaining Homeostasis of Brains. J. Pharmacol. Sci. 2021, 145 (3), 223–227. 10.1016/j.jphs.2020.12.007.

(10) Molina-Gonzalez, I.; Holloway, R. K.; Jiwaji, Z.; Dando, O.; Kent, S. A.; Emelianova, K.; Lloyd, A. F.; Forbes, L. H.; Mahmood, A.; Skripuletz, T.; Gudi, V.; Febery, J. A.; Johnson, J. A.; Fowler, J. H.; Kuhlmann, T.; Williams, A.; Chandran, S.; Stangel, M.; Howden, A. J. M.; Hardingham, G. E.; Miron, V. E. Astrocyte-Oligodendrocyte Interaction Regulates Central Nervous System Regeneration. Nat. Commun. 2023, 14 (1), 3372. 10.1038/s41467-023-39046-8.

(11) Baldwin, K. T.; Murai, K. K.; Khakh, B. S. Astrocyte Morphology. Trends Cell Biol. 2024, 34 (7), 547–565. 10.1016/j.tcb.2023.09.006.

(12) Lattke, M.; Guillemot, F. Understanding Astrocyte Differentiation: Clinical Relevance, Technical Challenges, and New Opportunities in the Omics Era. Wires Mech. Dis. 2022, 14 (5), e1557. 10.1002/wsbm.1557.

(13) Sloan, S. A.; Barres, B. A. Mechanisms of Astrocyte Development and Their Contributions to Neurodevelopmental Disorders. Curr. Opin. Neurobiol. 2014, 27, 75–81. 10.1016/j.conb.2014.03.005.

(14) Stipursky, J.; Francis, D.; Dezonne, R. S.; Bérgamo de Araújo, A. P.; Souza, L.; Moraes, C. A.; Alcantara Gomes, F. C. TGF-Β1 Promotes Cerebral Cortex Radial Glia-Astrocyte Differentiation in Vivo. Front. Cell. Neurosci. 2014, 8, 393. 10.3389/fncel.2014.00393.

(15) Cahoy, J. D.; Emery, B.; Kaushal, A.; Foo, L. C.; Zamanian, J. L.; Christopherson, K. S.; Xing, Y.; Lubischer, J. L.; Krieg, P. A.; Krupenko, S. A.; Thompson, W. J.; Barres, B. A. A Transcriptome Database for Astrocytes, Neurons, and Oligodendrocytes: A New Resource for Understanding Brain Development and Function. J. Neurosci. Off. J. Soc. Neurosci. 2008, 28 (1), 264–278. 10.1523/JNEUROSCI.4178-07.2008.

(16) van Meer, G.; Voelker, D. R.; Feigenson, G. W. Membrane Lipids: Where They Are and How They Behave. Nat. Rev. Mol. Cell Biol. 2008, 9 (2), 112–124. 10.1038/nrm2330.

(17) Gault, C. R.; Obeid, L. M.; Hannun, Y. A. An Overview of Sphingolipid Metabolism: From Synthesis to Breakdown. Adv. Exp. Med. Biol. 2010, 688, 1. 10.1007/978-1-4419-6741-1_1.

(18) Simons, K.; Toomre, D. Lipid Rafts and Signal Transduction. Nat. Rev. Mol. Cell Biol. 2000, 1 (1), 31–39. 10.1038/35036052.

(19) Maceyka, M.; Spiegel, S. Sphingolipid Metabolites in Inflammatory Disease. Nature 2014, 510 (7503), 58–67. 10.1038/nature13475.

(20) Sonnino, S.; Prinetti, A. The Role of Sphingolipids in Neuronal Plasticity of the Brain. J. Neurochem. 2016, 137 (4), 485–488. 10.1111/jnc.13589.

(21) Yu, R. K.; Nakatani, Y.; Yanagisawa, M. The Role of Glycosphingolipid Metabolism in the Developing Brain. J. Lipid Res. 2009, 50 (SUPPL.), 440–445. 10.1194/jlr.R800028-JLR200.

(22) Bassi, R.; Anelli, V.; Giussani, P.; Tettamanti, G.; Viani, P.; Riboni, L. Sphingosine-1-Phosphate Is Released by Cerebellar Astrocytes in Response to bFGF and Induces Astrocyte Proliferation through Gi-Protein-Coupled Receptors. Glia 2006, 53 (6), 621–630. 10.1002/glia.20324.

(23) Sorensen, S. D.; Nicole, O.; Peavy, R. D.; Montoya, L. M.; Lee, C. J.; Murphy, T. J.; Traynelis, S. F.; Hepler, J. R. Common Signaling Pathways Link Activation of Murine PAR-1, LPA, and S1P Receptors to Proliferation of Astrocytes. Mol. Pharmacol. 2003, 64 (5), 1199–1209. 10.1124/mol.64.5.1199.

(24) Young, N.; Van Brocklyn, J. R. Roles of Sphingosine-1-Phosphate (S1P) Receptors in Malignant Behavior of Glioma Cells. Differential Effects of S1P2 on Cell Migration and Invasiveness. Exp. Cell Res. 2007, 313 (8), 1615–1627. 10.1016/j.yexcr.2007.02.009.

(25) Cencetti, F.; Bernacchioni, C.; Bruno, M.; Squecco, R.; Idrizaj, E.; Berbeglia, M.; Bruni, P.; Donati, C. Sphingosine 1-Phosphate-Mediated Activation of Ezrin-Radixin-Moesin Proteins Contributes to Cytoskeletal Remodeling and Changes of Membrane Properties in Epithelial Otic Vesicle Progenitors. Biochim. Biophys. Acta BBA - Mol. Cell Res. 2019, 1866 (4), 554–565. 10.1016/j.bbamcr.2018.12.007.

(26) Paik, J. H.; Chae, S.; Lee, M.-J.; Thangada, S.; Hla, T. Sphingosine 1-Phosphate-Induced Endothelial Cell Migration Requires the Expression of EDG-1 and EDG-3 Receptors and Rho-Dependent Activation of Αvβ3- and Β1-Containing Integrins *. J. Biol. Chem. 2001, 276 (15), 11830–11837. 10.1074/jbc.M009422200.

(27) Toman, R. E.; Payne, S. G.; Watterson, K. R.; Maceyka, M.; Lee, N. H.; Milstien, S.; Bigbee, J. W.; Spiegel, S. Differential Transactivation of Sphingosine-1-Phosphate Receptors Modulates NGF-Induced Neurite Extension. J. Cell Biol. 2004, 166 (3), 381–392. 10.1083/jcb.200402016.

(28) Singh, S. K.; Kordula, T.; Spiegel, S. Neuronal Contact Upregulates Astrocytic Sphingosine-1-Phosphate Receptor 1 to Coordinate Astrocyte-Neuron Cross Communication. Glia 2022, 70 (4), 712–727. 10.1002/glia.24135.

(29) Chen, J.; Stork, T.; Kang, Y.; Nardone, K. A. M.; Auer, F.; Farrell, R. J.; Jay, T. R.; Heo, D.; Sheehan, A.; Paton, C.; Nagel, K. I.; Schoppik, D.; Monk, K. R.; Freeman, M. R. Astrocyte Growth Is Driven by the Tre1/S1pr1 Phospholipid-Binding G Protein-Coupled Receptor. Neuron 2024, 112 (1), 93–112.e10. 10.1016/j.neuron.2023.11.008.

(30) Singh, S. K.; Stogsdill, J. A.; Pulimood, N. S.; Dingsdale, H.; Kim, Y. H.; Pilaz, L.-J.; Kim, I. H.; Manhaes, A. C.; Rodrigues, W. S.; Pamukcu, A.; Enustun, E.; Ertuz, Z.; Scheiffele, P.; Soderling, S. H.; Silver, D. L.; Ji, R.-R.; Medina, A. E.; Eroglu, C. Astrocytes Assemble Thalamocortical Synapses by Bridging NRX1α and NL1 via Hevin. Cell 2016, 164 (1), 183–196. 10.1016/j.cell.2015.11.034.

(31) Uezu, A.; Kanak, D. J.; Bradshaw, T. W. A.; Soderblom, E. J.; Catavero, C. M.; Burette, A. C.; Weinberg, R. J.; Soderling, S. H. Identification of an Elaborate Complex Mediating Postsynaptic Inhibition. Science 2016, 353 (6304), 1123–1129. 10.1126/science.aag0821.

(32) Liddelow, S. A. Modern Approaches to Investigating Non-Neuronal Aspects of Alzheimer’s Disease. FASEB J. Off. Publ. Fed. Am. Soc. Exp. Biol. 2019, 33 (2), 1528–1535. 10.1096/fj.201802592.

(33) Gonzalez-Cabrera, P. J.; Jo, E.; Sanna, M. G.; Brown, S.; Leaf, N.; Marsolais, D.; Schaeffer, M.-T.; Chapman, J.; Cameron, M.; Guerrero, M.; Roberts, E.; Rosen, H. Full Pharmacological Efficacy of a Novel S1P1 Agonist That Does Not Require S1P-like Headgroup Interactions. Mol. Pharmacol. 2008, 74 (5), 1308–1318. 10.1124/mol.108.049783.

(34) Acaz-Fonseca, E.; Ortiz-Rodriguez, A.; Azcoitia, I.; Garcia-Segura, L. M.; Arevalo, M.-A. Notch Signaling in Astrocytes Mediates Their Morphological Response to an Inflammatory Challenge. Cell Death Discov. 2019, 5 (1), 85. 10.1038/s41420-019-0166-6.

(35) Xie, Y.; Kuan, A. T.; Wang, W.; Herbert, Z. T.; Mosto, O.; Olukoya, O.; Adam, M.; Vu, S.; Kim, M.; Tran, D.; Gómez, N.; Charpentier, C.; Sorour, I.; Lacey, T. E.; Tolstorukov, M. Y.; Sabatini, B. L.; Lee, W.-C. A.; Harwell, C. C. Astrocyte-Neuron Crosstalk through Hedgehog Signaling Mediates Cortical Synapse Development. Cell Rep. 2022, 38 (8), 110416. 10.1016/j.celrep.2022.110416.

(36) Hong, S.; Song, M.-R. STAT3 but Not STAT1 Is Required for Astrocyte Differentiation. PLOS ONE 2014, 9 (1), e86851. 10.1371/journal.pone.0086851.

(37) Yanagida, K.; Liu, C. H.; Faraco, G.; Galvani, S.; Smith, H. K.; Burg, N.; Anrather, J.; Sanchez, T.; Iadecola, C.; Hla, T. Size-Selective Opening of the Blood–Brain Barrier by Targeting Endothelial Sphingosine 1–Phosphate Receptor 1. Proc. Natl. Acad. Sci. 2017, 114 (17), 4531–4536. 10.1073/pnas.1618659114.

(38) Baldwin, K. T.; Tan, C. X.; Strader, S. T.; Jiang, C.; Savage, J. T.; Elorza-Vidal, X.; Contreras, X.; Rülicke, T.; Hippenmeyer, S.; Estévez, R.; Ji, R. R.; Eroglu, C. HepaCAM Controls Astrocyte Self-Organization and Coupling. Neuron 2021, 109 (15), 2427–2442.e10. 10.1016/j.neuron.2021.05.025.

(39) Lanjakornsiripan, D.; Pior, B. J.; Kawaguchi, D.; Furutachi, S.; Tahara, T.; Katsuyama, Y.; Suzuki, Y.; Fukazawa, Y.; Gotoh, Y. Layer-Specific Morphological and Molecular Differences in Neocortical Astrocytes and Their Dependence on Neuronal Layers. Nat. Commun. 2018, 9 (1). 10.1038/s41467-018-03940-3.

(40) Singh, S. K.; Weigel, C.; Brown, R. D. R.; Green, C. D.; Tuck, C.; Salvemini, D.; Spiegel, S. FTY720/Fingolimod Mitigates Paclitaxel-Induced Sparcl1-Driven Neuropathic Pain and Breast Cancer Progression. FASEB J. Off. Publ. Fed. Am. Soc. Exp. Biol. 2024, 38 (15), e23872. 10.1096/fj.202401277R.

(41) Chen, Z.; Doyle, T. M.; Luongo, L.; Largent-Milnes, T. M.; Giancotti, L. A.; Kolar, G.; Squillace, S.; Boccella, S.; Walker, J. K.; Pendleton, A.; Spiegel, S.; Neumann, W. L.; Vanderah, T. W.; Salvemini, D. Sphingosine-1-Phosphate Receptor 1 Activation in Astrocytes Contributes to Neuropathic Pain. Proc. Natl. Acad. Sci. U. S. A. 2019, 116 (21), 10557–10562. 10.1073/pnas.1820466116.

(42) Kim, S. K.; Hayashi, H.; Ishikawa, T.; Shibata, K.; Shigetomi, E.; Shinozaki, Y.; Inada, H.; Roh, S. E.; Kim, S. J.; Lee, G.; Bae, H.; Moorhouse, A. J.; Mikoshiba, K.; Fukazawa, Y.; Koizumi, S.; Nabekura, J. Cortical Astrocytes Rewire Somatosensory Cortical Circuits for Peripheral Neuropathic Pain. J. Clin. Invest. 2016, 126 (5), 1983–1997. 10.1172/JCI82859.

(43) Reeves, A. M. B.; Shigetomi, E.; Khakh, B. S. Bulk Loading of Calcium Indicator Dyes to Study Astrocyte Physiology: Key Limitations and Improvements Using Morphological Maps. J. Neurosci. 2011, 31 (25), 9353–9358. 10.1523/JNEUROSCI.0127-11.2011.

(44) Batiuk, M. Y.; Martirosyan, A.; Wahis, J.; de Vin, F.; Marneffe, C.; Kusserow, C.; Koeppen, J.; Viana, J. F.; Oliveira, J. F.; Voet, T.; Ponting, C. P.; Belgard, T. G.; Holt, M. G. Identification of Region-Specific Astrocyte Subtypes at Single Cell Resolution. Nat. Commun. 2020, 11 (1), 1220. 10.1038/s41467-019-14198-8.

(45) Peng, H.; Xie, P.; Liu, L.; Kuang, X.; Wang, Y.; Qu, L.; Gong, H.; Jiang, S.; Li, A.; Ruan, Z.; Ding, L.; Yao, Z.; Chen, C.; Chen, M.; Daigle, T. L.; Dalley, R.; Ding, Z.; Duan, Y.; Feiner, A.; He, P.; Hill, C.; Hirokawa, K. E.; Hong, G.; Huang, L.; Kebede, S.; Kuo, H. C.; Larsen, R.; Lesnar, P.; Li, L.; Li, Q.; Li, X.; Li, Y.; Li, Y.; Liu, A.; Lu, D.; Mok, S.; Ng, L.; Nguyen, T. N.; Ouyang, Q.; Pan, J.; Shen, E.; Song, Y.; Sunkin, S. M.; Tasic, B.; Veldman, M. B.; Wakeman, W.; Wan, W.; Wang, P.; Wang, Q.; Wang, T.; Wang, Y.; Xiong, F.; Xiong, W.; Xu, W.; Ye, M.; Yin, L.; Yu, Y.; Yuan, J.; Yuan, J.; Yun, Z.; Zeng, S.; Zhang, S.; Zhao, S.; Zhao, Z.; Zhou, Z.; Huang, Z. J.; Esposito, L.; Hawrylycz, M. J.; Sorensen, S. A.; Yang, X. W.; Zheng, Y.; Gu, Z.; Xie, W.; Koch, C.; Luo, Q.; Harris, J. A.; Wang, Y.; Zeng, H. Morphological Diversity of Single Neurons in Molecularly Defined Cell Types. Nature 2021, 598 (7879), 174–181. 10.1038/s41586-021-03941-1.

(46) Tan, C. X.; Bindu, D. S.; Hardin, E. J.; Sakers, K.; Baumert, R.; Ramirez, J. J.; Savage, J. T.; Eroglu, C. δ-Catenin Controls Astrocyte Morphogenesis via Layer-Specific Astrocyte–Neuron Cadherin Interactions. J. Cell Biol. 2023, 222 (11), e202303138. 10.1083/jcb.202303138.

(47) Yan, X.; Li, S.; Wang, J.; Wang, Q.; Xie, X.; Li, M. Sphingosine-1-Phosphate Induces Pulmonary Artery Smooth Muscle Cell Proliferation, Migration and Pulmonary Arterial Remodeling by Modulating Sonic Hedgehog Signaling Effector FoxM1. Chin. Med. J. (Engl.) 2026. 10.1097/CM9.0000000000004041.

(48) Bayraktar, O. A.; Bartels, T.; Holmqvist, S.; Kleshchevnikov, V.; Martirosyan, A.; Polioudakis, D.; Ben Haim, L.; Young, A. M. H.; Batiuk, M. Y.; Prakash, K.; Brown, A.; Roberts, K.; Paredes, M. F.; Kawaguchi, R.; Stockley, J. H.; Sabeur, K.; Chang, S. M.; Huang, E.; Hutchinson, P.; Ullian, E. M.; Hemberg, M.; Coppola, G.; Holt, M. G.; Geschwind, D. H.; Rowitch, D. H. Astrocyte Layers in the Mammalian Cerebral Cortex Revealed by a Single-Cell in Situ Transcriptomic Map. Nat. Neurosci. 2020, 23 (4), 500– 509. 10.1038/s41593-020-0602-1.

(49) Lee, J. Y. Normal and Disordered Formation of the Cerebral Cortex : Normal Embryology, Related Molecules, Types of Migration, Migration Disorders. J. Korean Neurosurg. Soc. 2019, 62 (3), 265–271. 10.3340/jkns.2019.0098.

(50) Wu, M. W.; Kourdougli, N.; Portera-Cailliau, C. Network State Transitions during Cortical Development. Nat. Rev. Neurosci. 2024, 25 (8), 535–552. 10.1038/s41583-024-00824-y.

(51) Bocchi, R.; Thorwirth, M.; Simon-Ebert, T.; Koupourtidou, C.; Clavreul, S.; Kolf, K.; Della Vecchia, P.; Bottes, S.; Jessberger, S.; Zhou, J.; Wani, G.; Pilz, G.-A.; Ninkovic, J.; Buffo, A.; Sirko, S.; Götz, M.; Fischer-Sternjak, J. Astrocyte Heterogeneity Reveals Region-Specific Astrogenesis in the White Matter. Nat. Neurosci. 2025, 28 (3), 457–469. 10.1038/s41593-025-01878-6.

(52) Cheng, Y.-T.; Luna-Figueroa, E.; Woo, J.; Chen, H.-C.; Lee, Z.-F.; Harmanci, A. S.; Deneen, B. Inhibitory Input Directs Astrocyte Morphogenesis through Glial GABABR. Nature 2023, 617 (7960), 369–376. 10.1038/s41586-023-06010-x.

(53) Holt, L. M.; Hernandez, R. D.; Pacheco, N. L.; Torres Ceja, B.; Hossain, M.; Olsen, M. L. Astrocyte Morphogenesis Is Dependent on BDNF Signaling via Astrocytic TrkB.T1. eLife 2019, 8, 1–27. 10.7554/eLife.44667.

(54) Bushong, E. A.; Martone, M. E.; Jones, Y. Z.; Ellisman, M. H. Protoplasmic Astrocytes in CA1 Stratum Radiatum Occupy Separate Anatomical Domains. J. Neurosci. 2002, 22 (1), 183–192. 10.1523/jneurosci.22-01-00183.2002.

(55) Shigetomi, E.; Bushong, E. A.; Haustein, M. D.; Tong, X.; Jackson-Weaver, O.; Kracun, S.; Xu, J.; Sofroniew, M. V.; Ellisman, M. H.; Khakh, B. S. Imaging Calcium Microdomains within Entire Astrocyte Territories and Endfeet with GCaMPs Expressed Using Adeno-Associated Viruses. J. Gen. Physiol. 2013, 141 (5), 633–647. 10.1085/jgp.201210949.

(56) Hayashi, M. K.; Sato, K.; Sekino, Y. Neurons Induce Tiled Astrocytes with Branches That Avoid Each Other. Int. J. Mol. Sci. 2022, 23 (8). 10.3390/ijms23084161.

(57) Sansom, S. N.; Livesey, F. J. Gradients in the Brain: The Control of the Development of Form and Function in the Cerebral Cortex. Cold Spring Harb. Perspect. Biol. 2009, 1 (2), a002519. 10.1101/cshperspect.a002519.

(58) Neal, M.; Luo, D.; Harischandra, D. S.; Gordon, R.; Sarkar, S.; Jin, H.; Anantharam, V.; Désaubry, L.; Kanthasamy, A.; Kanthasamy, A. Prokineticin-2 Promotes Chemotaxis and Alternative A2 Reactivity of Astrocytes. Glia 2018, 66 (10), 2137–2157. 10.1002/glia.23467.

(59) Bicknell, B. A.; Pujic, Z.; Feldner, J.; Vetter, I.; Goodhill, G. J. Chemotactic Responses of Growing Neurites to Precisely Controlled Gradients of Nerve Growth Factor. Sci. Data 2018, 5 (1), 180183. 10.1038/sdata.2018.183.

(60) Rivera, J.; Proia, R. L.; Olivera, A. THE ALLIANCE OF SPHINGOSINE-1-PHOSPHATE AND ITS RECEPTORS IN IMMUNITY. Nat. Rev. Immunol. 2008, 8 (10), 753–763. 10.1038/nri2400.

(61) Kharel, Y.; Huang, T.; Salamon, A.; Harris, T. E.; Santos, W. L.; Lynch, K. R. Mechanism of Sphingosine 1-Phosphate Clearance from Blood. Biochem. J. 2020, 477 (5), 925–935. 10.1042/BCJ20190730.

(62) Kono, M.; Mi, Y.; Liu, Y.; Sasaki, T.; Allende, M. L.; Wu, Y.-P.; Yamashita, T.; Proia, R. L. The Sphingosine-1-Phosphate Receptors S1P1, S1P2, and S1P3 Function Coordinately during Embryonic Angiogenesis*. J. Biol. Chem. 2004, 279 (28), 29367–29373. 10.1074/jbc.M403937200.

(63) Gatfield, J.; Monnier, L.; Studer, R.; Bolli, M. H.; Steiner, B.; Nayler, O. Sphingosine-1-Phosphate (S1P) Displays Sustained S1P1 Receptor Agonism and Signaling through S1P Lyase-Dependent Receptor Recycling. Cell. Signal. 2014, 26 (7), 1576–1588. 10.1016/j.cellsig.2014.03.029.

(64) Yuan, Y.; Jia, G.; Wu, C.; Wang, W.; Cheng, L.; Li, Q.; Li, Z.; Luo, K.; Yang, S.; Yan, W.; Su, Z.; Shao, Z. Structures of Signaling Complexes of Lipid Receptors S1PR1 and S1PR5 Reveal Mechanisms of Activation and Drug Recognition. Cell Res. 2021, 31 (12), 1263– 1274. 10.1038/s41422-021-00566-x.

(65) Donati, C.; Bruni, P. Sphingosine 1-Phosphate Regulates Cytoskeleton Dynamics: Implications in Its Biological Response. Biochim. Biophys. Acta BBA - Biomembr. 2006, 1758 (12), 2037–2048. 10.1016/j.bbamem.2006.06.015.

(66) Ko, P.; Kim, D.; You, E.; Jung, J.; Oh, S.; Kim, J.; Lee, K.-H.; Rhee, S. Extracellular Matrix Rigidity-Dependent Sphingosine-1-Phosphate Secretion Regulates Metastatic Cancer Cell Invasion and Adhesion. Sci. Rep. 2016, 6 (1), 21564. 10.1038/srep21564.

(67) Lee, J.-F.; Ozaki, H.; Zhan, X.; Wang, E.; Hla, T.; Lee, M.-J. Sphingosine-1-Phosphate Signaling Regulates Lamellipodia Localization of Cortactin Complexes in Endothelial Cells. Histochem. Cell Biol. 2006, 126 (3), 297–304. 10.1007/s00418-006-0143-z.

(68) Sassoli, C.; Pierucci, F.; Zecchi-Orlandini, S.; Meacci, E. Sphingosine 1-Phosphate (S1P)/ S1P Receptor Signaling and Mechanotransduction: Implications for Intrinsic Tissue Repair/Regeneration. Int. J. Mol. Sci. 2019, 20 (22), 5545. 10.3390/ijms20225545.

(69) Haber, M.; Zhou, L.; Murai, K. K. Cooperative Astrocyte and Dendritic Spine Dynamics at Hippocampal Excitatory Synapses. J. Neurosci. Off. J. Soc. Neurosci. 2006, 26 (35), 8881– 8891. 10.1523/JNEUROSCI.1302-06.2006.

(70) Murk, K.; Blanco Suarez, E. M.; Cockbill, L. M. R.; Banks, P.; Hanley, J. G. The Antagonistic Modulation of Arp2/3 Activity by N-WASP, WAVE2 and PICK1 Defines Dynamic Changes in Astrocyte Morphology. J. Cell Sci. 2013, 126 (Pt 17), 3873–3883. 10.1242/jcs.125146.

(71) Hobson, J. P.; Rosenfeldt, H. M.; Barak, L. S.; Olivera, A.; Poulton, S.; Caron, M. G.; Milstien, S.; Spiegel, S. Role of the Sphingosine-1-Phosphate Receptor EDG-1 in PDGF-Induced Cell Motility. Science 2001, 291 (5509), 1800–1803. 10.1126/science.1057559.

(72) Lee, M.-J.; Thangada, S.; Paik, J.-H.; Sapkota, G. P.; Ancellin, N.; Chae, S.-S.; Wu, M.; Morales-Ruiz, M.; Sessa, W. C.; Alessi, D. R.; Hla, T. Akt-Mediated Phosphorylation of the G Protein-Coupled Receptor EDG-1 Is Required for Endothelial Cell Chemotaxis. Mol. Cell 2001, 8 (3), 693–704. 10.1016/S1097-2765(01)00324-0.

(73) Schiweck, J.; Eickholt, B. J.; Murk, K. Important Shapeshifter: Mechanisms Allowing Astrocytes to Respond to the Changing Nervous System during Development, Injury and Disease. Front. Cell. Neurosci. 2018, 12 (August), 1–17. 10.3389/fncel.2018.00261.

(74) Schober, A. L.; Wicki-Stordeur, L. E.; Murai, K. K.; Swayne, L. A. Foundations and Implications of Astrocyte Heterogeneity during Brain Development and Disease. Trends Neurosci. 2022, 45 (9), 692–703. 10.1016/j.tins.2022.06.009.

(75) Badia-Soteras, A.; Mak, A.; Blok, T. M.; Boers-Escuder, C.; van den Oever, M. C.; Min, R.; Smit, A. B.; Verheijen, M. H. G. Astrocyte-Synapse Structural Plasticity in Neurodegenerative and Neuropsychiatric Diseases. Biol. Psychiatry 2025. 10.1016/j.biopsych.2025.04.011.

(76) Endo, F.; Kasai, A.; Soto, J. S.; Yu, X.; Qu, Z.; Hashimoto, H.; Gradinaru, V.; Kawaguchi, R.; Khakh, B. S. Molecular Basis of Astrocyte Diversity and Morphology across the CNS in Health and Disease. Science 2022, 378 (6619), eadc9020. 10.1126/science.adc9020.

