## Supplementary figures and images for "Sphingosine-1-Phosphate Receptor 1 regulates competition dependent astrocyte morphogenesis and tiling in murine cortex"

### S1

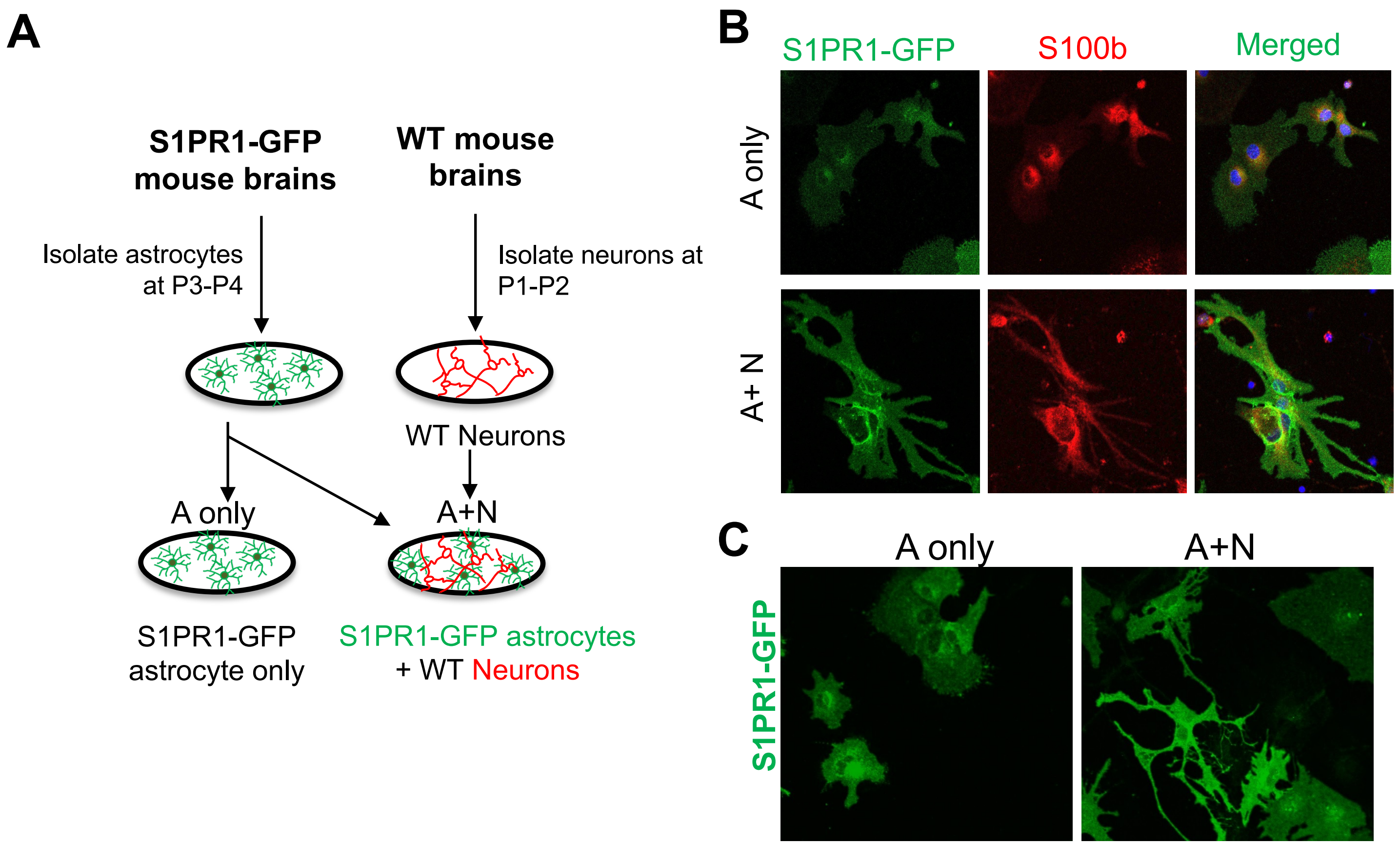

### S2

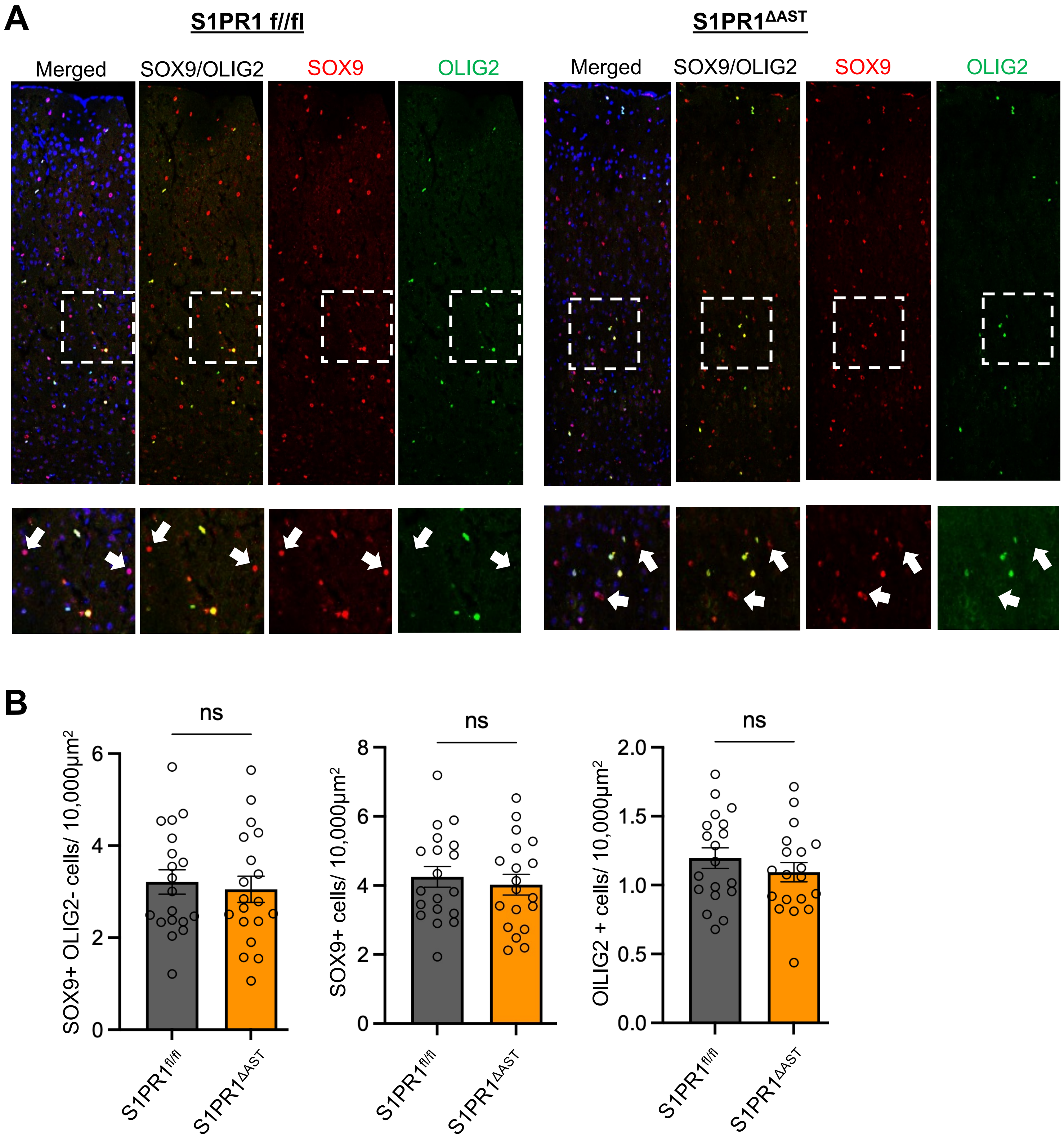

### S3

#### Fig. S3

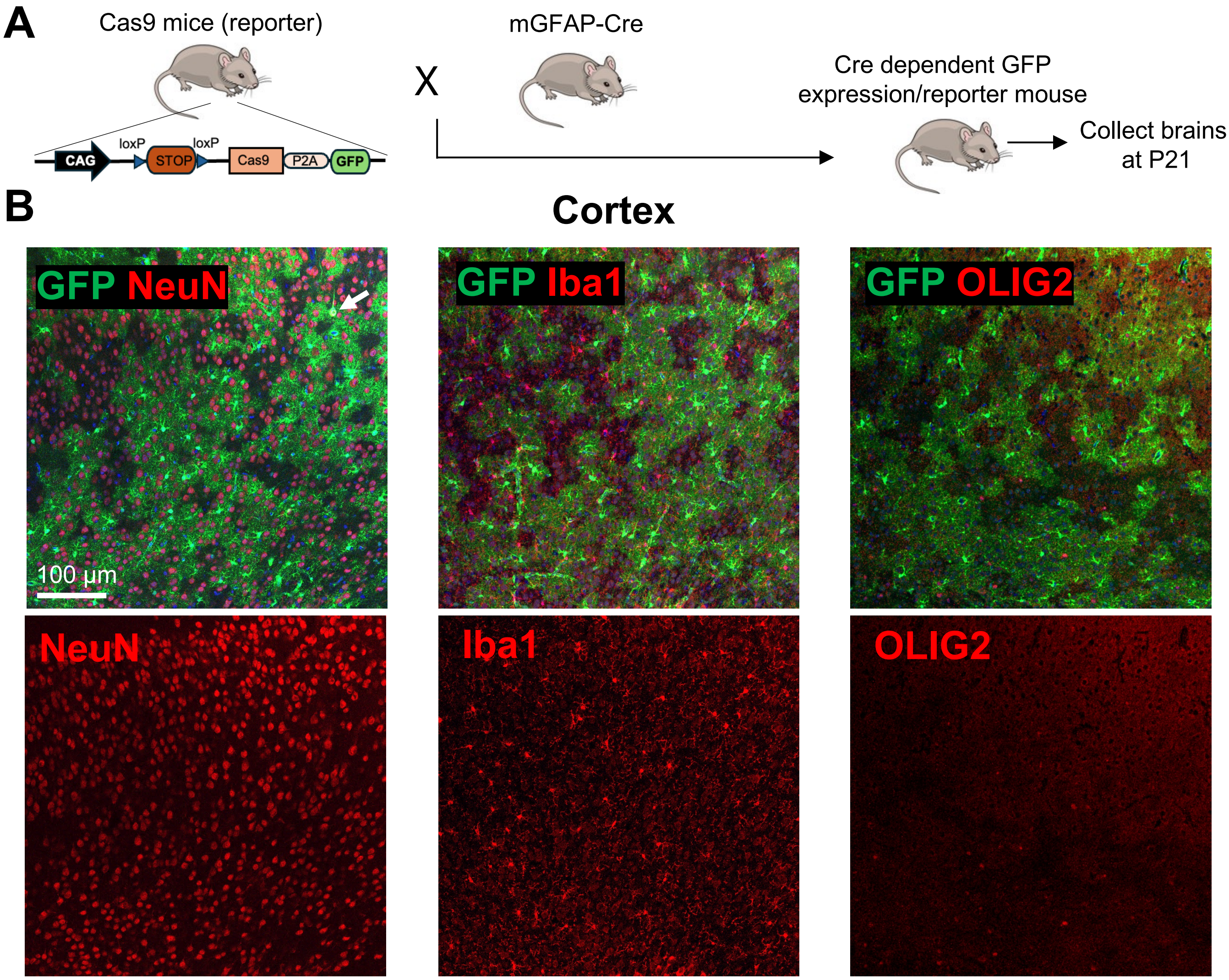


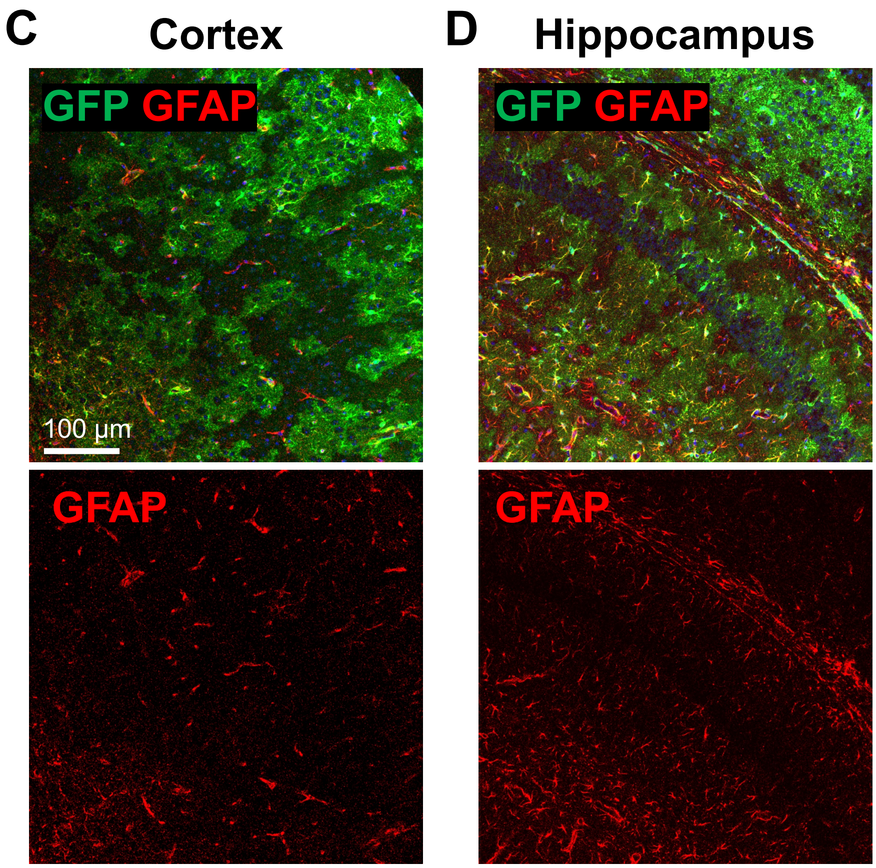


**E**

| Cell marker | % of GFP+ cells |
|---|---|
| NeuN+GFP+ (Neuron) | 0.63 ± 0.271 |
| Iba1+GFP+ (Microglia) | 0.00 ± 0.000 |
| OLIG2+GFP+ (Oligos) | 2.34 ± 0.868 |

### S4

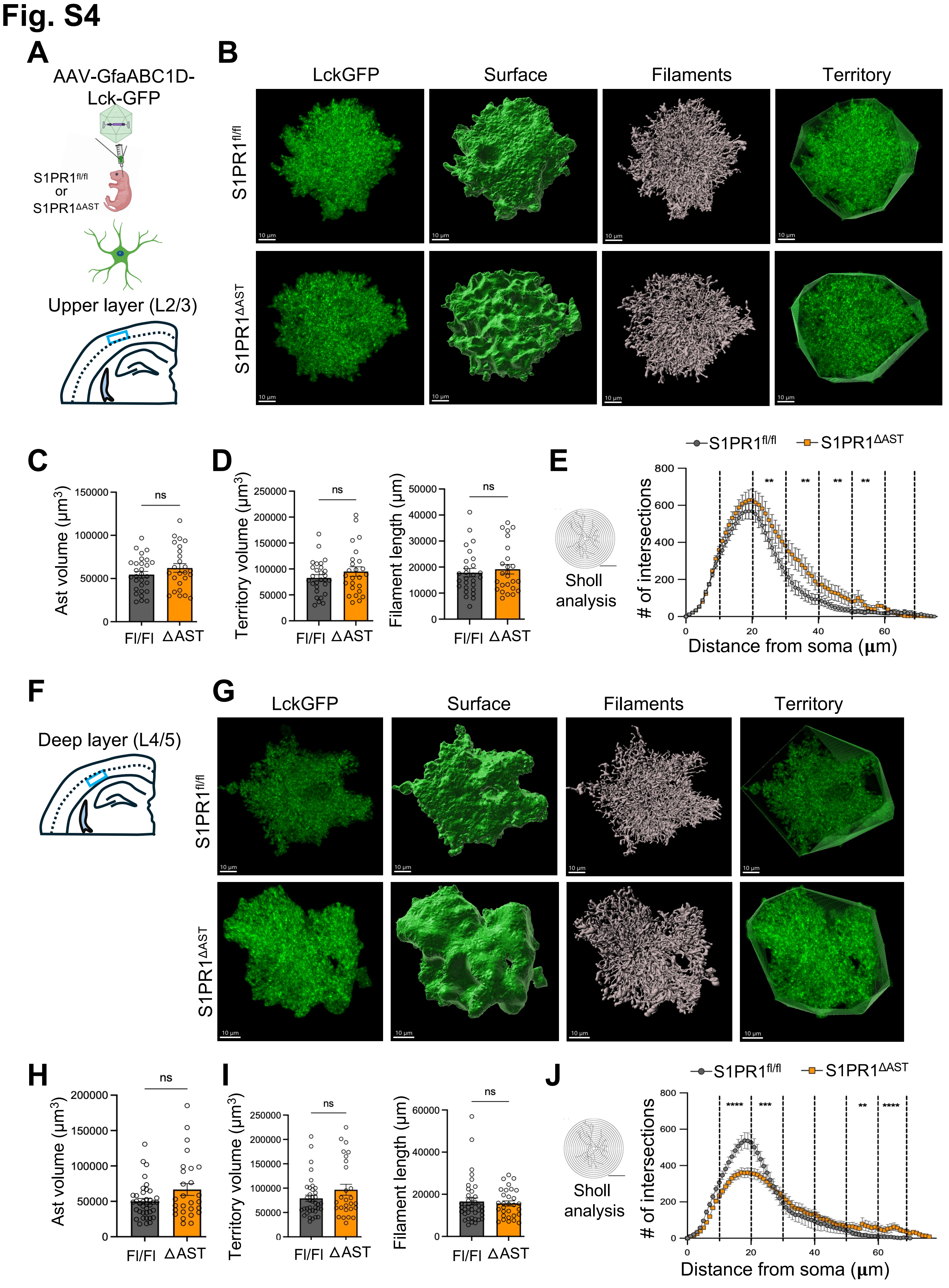

### S5

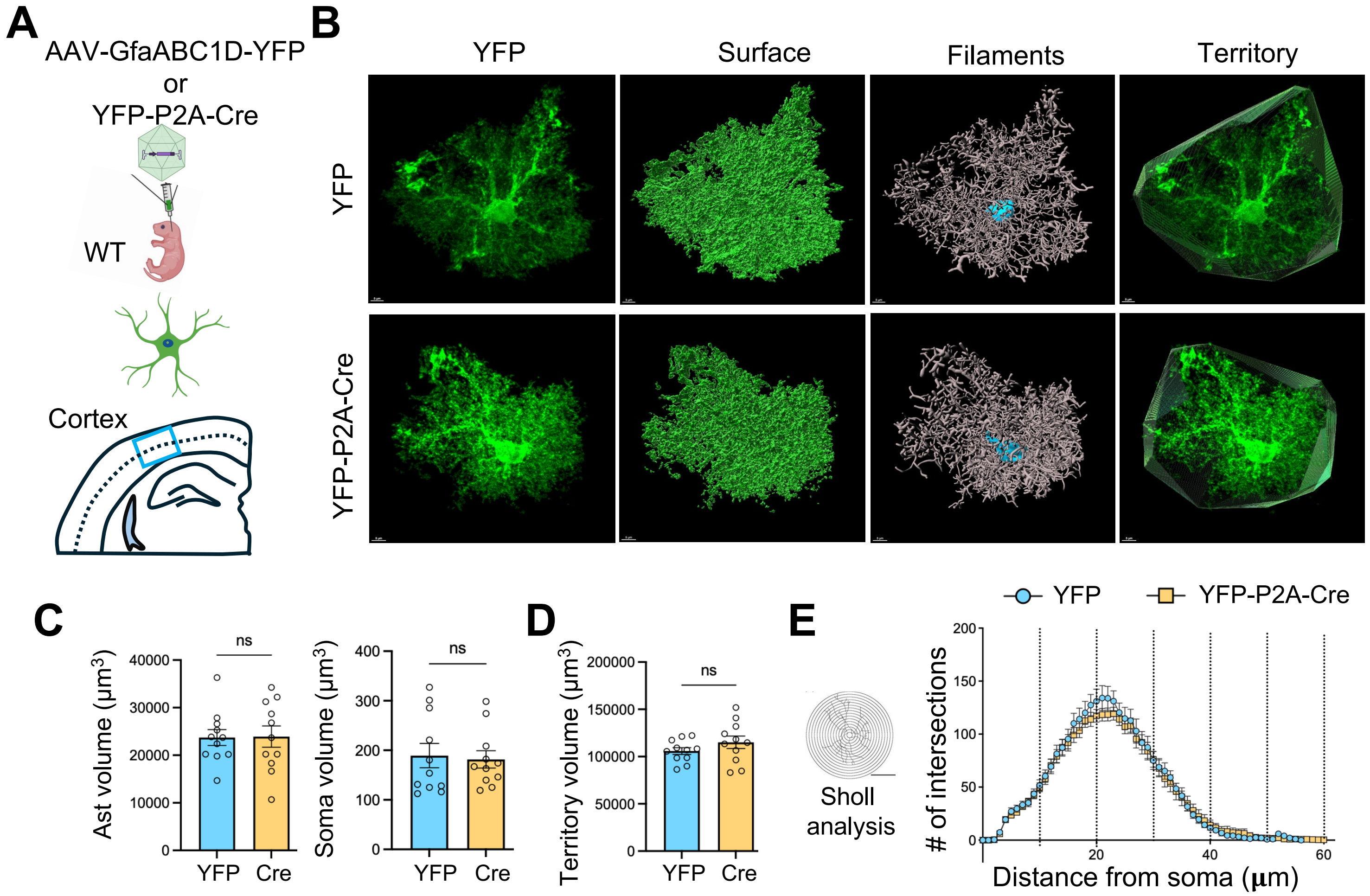
